# Modelling bistable tumour population dynamics to design effective treatment strategies

**DOI:** 10.1101/381418

**Authors:** Andrei R. Akhmetzhanov, Jong Wook Kim, Ryan Sullivan, Robert A. Beckman, Pablo Tamayo, Chen-Hsiang Yeang

## Abstract

Despite recent advances in targeted drugs and immunotherapy, cancer remains “the emperor of all maladies” due to inevitable emergence of resistance. Drug resistance is thought to be driven by mutations and/or dynamic plasticity that deregulate pathway activities and regulatory programs of a highly heterogeneous tumour. In this study, we propose a modelling framework to simulate population dynamics of heterogeneous tumour cells with reversible drug resistance. Drug sensitivity of a tumour cell is determined by its internal states, which are demarcated by coordinated activities of multiple interconnected oncogenic pathways. Transitions between cellular states depend on the effects of targeted drugs and regulatory relations between the pathways. Under this framework, we build a simple model to capture drug resistance characteristics of BRAF-mutant melanoma, where two cell states are described by two mutually inhibitory – main and alternative – pathways. We assume that cells with an activated main pathway are proliferative yet sensitive to the BRAF inhibitor, and cells with an activated alternative pathway are quiescent but resistant to the drug. We describe a dynamical process of tumour growth under various drug regimens using the explicit solution of mean-field equations. Based on these solutions, we compare efficacy of three treatment strategies: static treatments with continuous and constant dosages, periodic treatments with regular intermittent phases and drug holidays, and treatments derived from optimal control theory (OCT). Based on these analysis, periodic treatments outperform static treatments with a considerable margin, while treatments based on OCT outperform the best periodic treatment. Our results provide insights regarding optimal cancer treatment modalities for heterogeneous tumours, and may guide the development of optimal therapeutic strategies to circumvent drug resistance and due to tumour plasticity.

## Introduction

Despite the recent advances of targeted treatments and immunotherapy, complete cure of cancer is still rare due to the almost inevitable emergence of resistance [1,2]. Drug resistance arises from a wide range of complex processes at multiple levels [3–5]. At the tumour level, drug response emerges primarily from population dynamics of cancer cells. The most well-known mechanism is clonal evolution [6,7]. A bulk tumour is often populated by a heterogeneous group of cancer cells with diverse mutational landscapes, epigenomic states, pathway activities and gene regulatory programs. Treatments induce differential fitness of subclones and consequently select for the most resistant ones. In addition to clonal evolution, treatments may also induce differential plasticity of tumour cells by shifting their pathway activities [8] and regulatory programs [9]. The major difference between these two processes pertains to reversibility of drug resistance. For clonal evolution, drug sensitivity of an individual cell is determined solely by its genetic landscape and thus remains invariant during its life span. Drug resistance of a subclone is thus an irreversible phenotype as a resistant subclone will rarely back-mutate to a sensitive one. For differential plasticity, drug sensitivity of an individual cell is a reversible dynamic state rather than a fixed phenotype. Both mechanisms are supported by numerous experimental evidence (e.g., for clonal evolution, [10]; for differential plasticity of tumour cells [8,11–14]). However, the latter process may account for drug resistance that can be reverted when the therapy is lifted [15,16].

There is a rich literature of mathematical models for tumour clonal evolution that undergoes treatments (e.g., [17–23]). In contrast, models of cellular plastic responses to treatments are relatively limited and recent (e.g., [24–26], see also reviews [27–29]). The ultimate purpose of those models is to quantitatively predict tumour’s drug responses and employ this information to design effective treatments. Previously, we proposed a unified framework encompassing both mathematical models of tumour population dynamics and treatment design [30]. We considered a simple evolution model involved in subclones with differential resistance of two drugs, and tested efficacy of six heuristic treatment strategies by simulating population dynamics with a large number of parameter combinations informed by literature and clinical experience. We further extended the work by three drug systems and generalizing treatment strategies that incorporate long-term prediction of tumour population composition [31].

An important missing piece in this unified framework is a mathematical model that tackles reversible drug responses of cancer cells. To fill this gap, here we propose a model to explore the population dynamics of cancer cells during or after treatment with targeted agents that produce reversible effects. The state of each cell is inferred from the activities of multiple inter-dependent pathways, whereas the fitness of each cell depends on its internal state and the external environment (i.e. drug dosage). Treatments alter the cellular state composition of the population by both facilitating the single-cell state transitions in certain directions and inhibiting proliferation of subpopulations with different efficiencies. To capture the essence of the considered phenomenon, we consider only a simple scenario in which there are two major populations with mutually antagonistic signalling pathways. The main pathway promotes cell proliferation more efficiently but is also sensitive to a therapeutic agent. The alternative pathway induces slow proliferation but is also resistant to the agent. Due to reversibility of the states, the treatment strategy aims to balance between controlling the tumour load and reducing the influence of resistant cells.

Despite its simplicity, this model represents reasonably well the switching behaviour of BRAF^V600E^ mutant melanomas treated with BRAF inhibitor (vemurafenib) as previously reported [32,33]. Melanoma is a frequently lethal form of skin cancer with incidence rates continuing to rise in many countries [34]. Approximately half of cases harbour a BRAF^V600^ mutation [35]. The resulting mutation leads to constitutive activation of a down-stream cascade of the mitogen activated protein (MAPK) pathway including MEK and ERK that promote proliferation of cancer cells. Treatment with single-agent BRAF inhibitor disrupts MAPK signalling and achieves remission but leads to relapse in 6.7 months on average [32]. As of 2014, the standard of care for BRAF^V600^ mutant melanoma is the combination of inhibitors of BRAF and MEK [36,37]. Still, resistance emerges through numerous genetic mechanisms [38–41] or phenotypic changes, such as switching from the suppressed MITF pathway to an alternative pathway involving activation of NFκB [13,14,42,43]. The latter process of switching between two major oncogenic programs is accompanied by physiological changes in cancer cells [44].

These characteristics allow us to abstract the problem and formulate a minimal dynamical model that involves interactions of two distinct pathways. We derive the master equations for the dynamics of tumour population size and state composition and find their analytic solutions with arbitrary drug regimens. Based on these solutions, we compare the performance of three treatment strategies on simulated data: i) static treatments with continuous and constant dosages, ii) periodic treatments with regular intermittent treatment days and drug holidays, and iii) the treatments that minimize the tumour size change after fixed time periods.

## Model and methods

### 1. Assumptions and concepts

We consider a general and abstract case where cancer cells are self-replicating entities possessing different internal states with different proliferative capacities, distinct sensitivities to treatments, and consequently, the size and composition of the entire tumour population. From this more general case, we study a particular instance in detail as outlined below. The following simplifying assumptions are introduced:

i. The mutational landscape of tumour cells does not acquire major driver events (“hallmarks of cancer” [45]) or new resistance mutations during the course of therapy. In fact this may not be the case and integration of models of genetic evolution of resistance and cellular plasticity is an important future step.
ii. The dominant subclone of tumour cells, which is the focus here, by default possesses elevated activities of the “main” pathway that render it highly proliferative.
iii. Proliferation can also be sustained by an “alternative” pathway with lower efficiency. The two pathways are mutually inhibitory, thus without external intervention the tumour population is dominated by cells with an active main pathway over those with an active alternative pathway.
iv. A targeted agent or targeted combination inhibits proliferation of cells with an active main pathway and concomitantly facilitates activity transitions from the main to the alternative pathway.

Fig 1A illustrates the conceptual framework of the model. This may be seen as an archetypical bistable state model that may cover a particular class of internal “wirings” of a cell.

**Fig 1:**
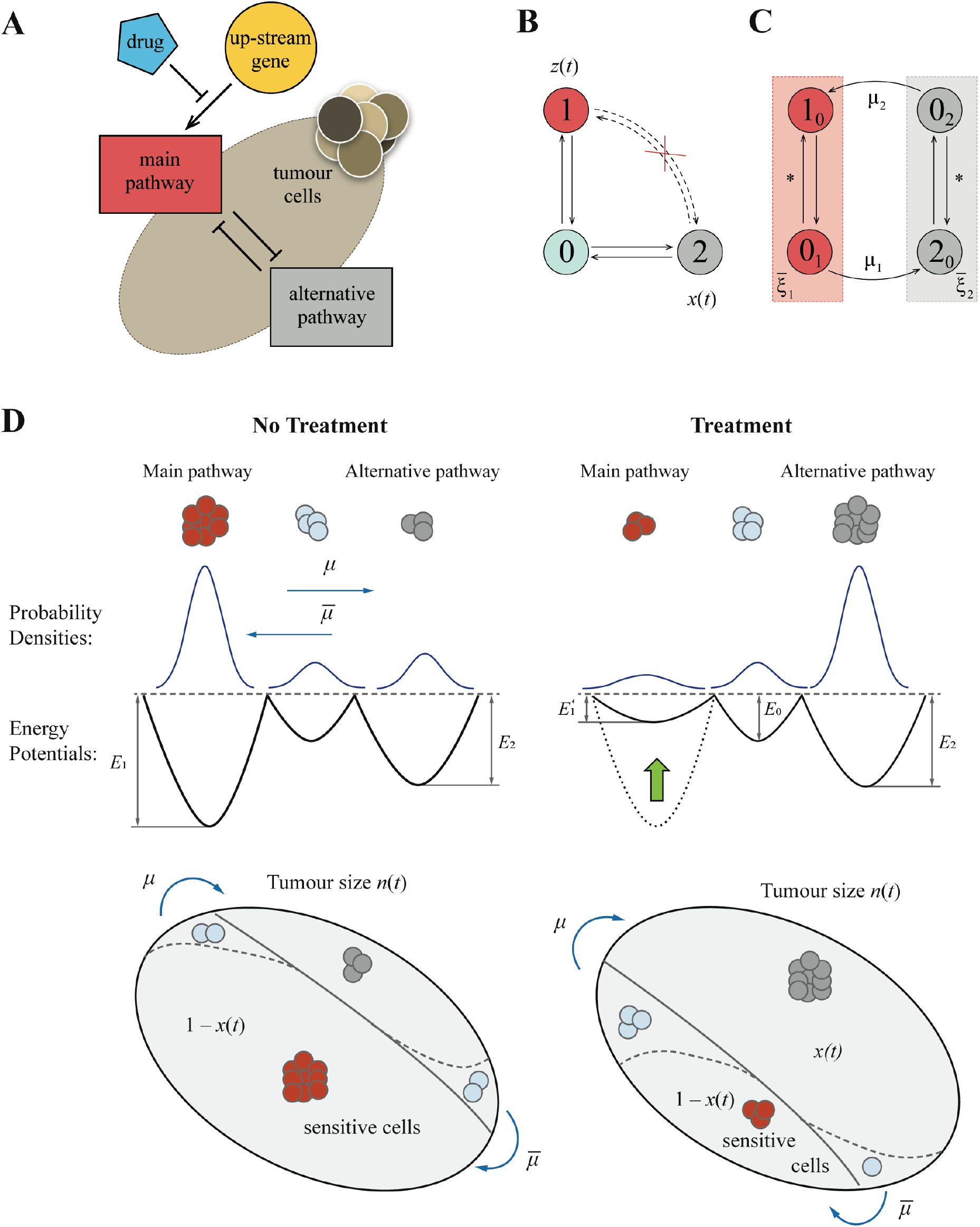
Mathematical modelling framework. (A) Schematic diagram of the pathway interaction within tumour cells. Activity of the main pathway is induced by an upstream gene, e.g. a driver oncogene, that can be blocked by the drug action. The regulation of the alternative resistant pathway remains unaffected. Both pathways are antagonistic to each other. (B) Original framework to model the switch between activated pathways “1” and “2” occurring through a transient state “0” where both pathways are shut down. (C) Flow diagram for a modified model with two states used to derive main equations. (D) Characteristics of a system without and with treatment. Each pathway activity is modelled by a stochastically moving particle in a potential well. The treatment changes only the potential for the main pathway, while the potential for the alternative pathway remains invariant.

Activities of the two pathways demarcate three cellular states (Fig 1BC): active main pathway and inactive alternative pathway (state “1”), inactive main pathway and active alternative pathway (state “2”), and inactive main and alternative pathways (state “0”). Simultaneous activation of both pathways is not allowed since they are mutually inhibitory. 0 is a transient state between 1 and 2.

Each single cell can encounter three stochastic events: proliferation, death, and state transition (1 → 0 → 2 or 2 → 0 → 1). The population dynamics of the birth-death process can be well approximated by ordinary differential equations. To determine the population dynamics of cellular states, we adopt a well-known approach from statistical physics by treating cells as particles undergoing Brownian motions inside a double-well potential [46]. In this model, each pathway possesses a double-well potential. The two equilibria of the system (the two local minima of the potential) represent up and down regulation of the two pathway activities (Fig 1D). Probabilities of staying in each state (and thereby the fraction of cells in each state) are determined by the “energy gap” between two local minima. Without external intervention, lower wells (more likely states) of the main pathway correspond to up-regulation of the main pathway and down-regulation of the alternative pathway.

The drug inhibits both activity of the main pathway and proliferation of main pathway-active cells, but has no effect on the alternative pathway. Mathematically it lifts the well of the up regulation state and lowers the well of the down regulation state of the main pathway potential function, and does not change the shape of the alternative pathway potential function. The extent of potential function change depends on administered dosage and consequently shifts the cellular state composition. We describe a magnitude of a relative shift by a variable σ that is constrained between zero and one. It decodes the treatment intensity with two extremalities: *σ* = 0 for no treatment, and *σ* =1 for a maximally tolerated dosage (MTD) administered. The latter shuts down the main pathway and turns on the alternative pathway, and thus drives drug resistance emergence.

This setting makes the disease incurable by continuous administration of a single therapeutic agent since the tumour inevitably relapses [47]. However, the patient’s life span can be significantly improved with proper arrangements of treatment dosage and schedule even of this single agent. We aim to minimize the tumour size after a fixed period of time. To fulfil this goal, an effective strategy should maintain a subtle balance of proliferative but sensitive cells vs. quiescent but resistant cells, such that the tumour is responsive to the drug but also has a limited growth rate. We compare the outcomes of three treatment strategies: a benchmark strategy of static treatment with a continuous and constant dosage, a heuristic strategy of periodic treatment with regular intermittent treatment days and drug holidays, and an analytic strategy derived from optimal control theory.

### 2. Dynamic equations

Population dynamics of cells follows a linear differential equation:

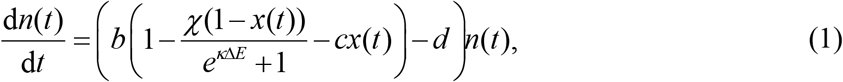

where *n*(*t*) denotes the total population size at time *t*, *x*(*t*) is the fraction of resistant cells (at state 2), *b* and *d* denote constant birth and death rates respectively. The fitness is penalized by two costs: the cost of resistance *c*, and the cost of inactivation of the main pathway *χ*. The latter is factored by the proportion of sensitive cells (1 − *x*(*t*)), multiplied by the fraction of them with currently inactivated main pathway ((*e*^*k*Δ*E*^ + 1)^−1^, where Δ*E* is the difference in depths of potential wells of the main pathway).

The complementary differential equation for the fraction of resistant cells *x*(*t*) is written by using the flow diagram in Fig 1D:

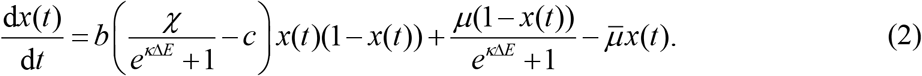

Here, the first term resembles the replicator dynamics [48], where the growth rate is equal to the difference between two fitness costs – it describes cell competition between different types. The two following terms indicate the transition flows between the sensitive state 1 and the resistant state 2. We denote the transition rates to and from the resistant states as *μ* and 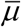 respectively.

We assume a linear dependence of the drug effect on the treatment intensity σ. Consequently, the double-well potential is defined by two quadratic polynomials that are attached to each other at the intermediate threshold point (Fig S1B). Then the values of *E*^−^ and Δ*E* in (1)–(2) are also quadratic functions of *σ*, see also Appendix A for more details.

We refer readers to Table 1 for summary of all model parameter and their estimates.

**Table 1.** Baseline parameter values used in this study.

| **Parameter** | **Variable** | **Value** | **Ref.** |
| --- | --- | --- | --- |
| Birth rate | $b$ | 0.14 day<sup>-1</sup> | [56] |
| Death rate | $d$ | 0.13 day<sup>-1</sup> | [56] |
| Cost of resistance | $c$ | 4% | |
| Cost of inactivation of both pathways | $\chi$ | 0.3 | |
| Characteristic switching time from the main to the alternative pathway | $\mu^{-1}$ | 28 days | |
| Characteristic switching time from the alternative to the main pathway | $\bar{\mu}^{-1}$ | 60 days | |
| Expression of the main pathway at down state | $\alpha$ | 0.3 | [57] |
| Threshold level for the production function of the main pathway | $\theta$ | 0.45 | [57] |
| Robustness parameter for the main pathway | $\kappa$ | 40 | |
| Effect of the drug on shifting the threshold level in activity of the main pathway | $L$ | 0.2 | |
| Initial level of resistance | $\varepsilon$ | 1% | [58] |

## Results

### 1. Static treatment

First, we report dynamics of tumour size of static treatments where the drug is administered at a constant dosage. The treatments with low intensities (e.g. *σ* = 0.2 in Fig 2A, red curve) yield exponential tumour growth. The treatments with intermediate or high intensities (e.g. *σ* ≥ 0.4 in Fig 2A, yellow and cyan curves) cause initial shrinkage of a tumour due to elimination of the proliferative cells, while the remaining resistant cells regrow and the tumour relapses in the later stage.

**Fig 2:**
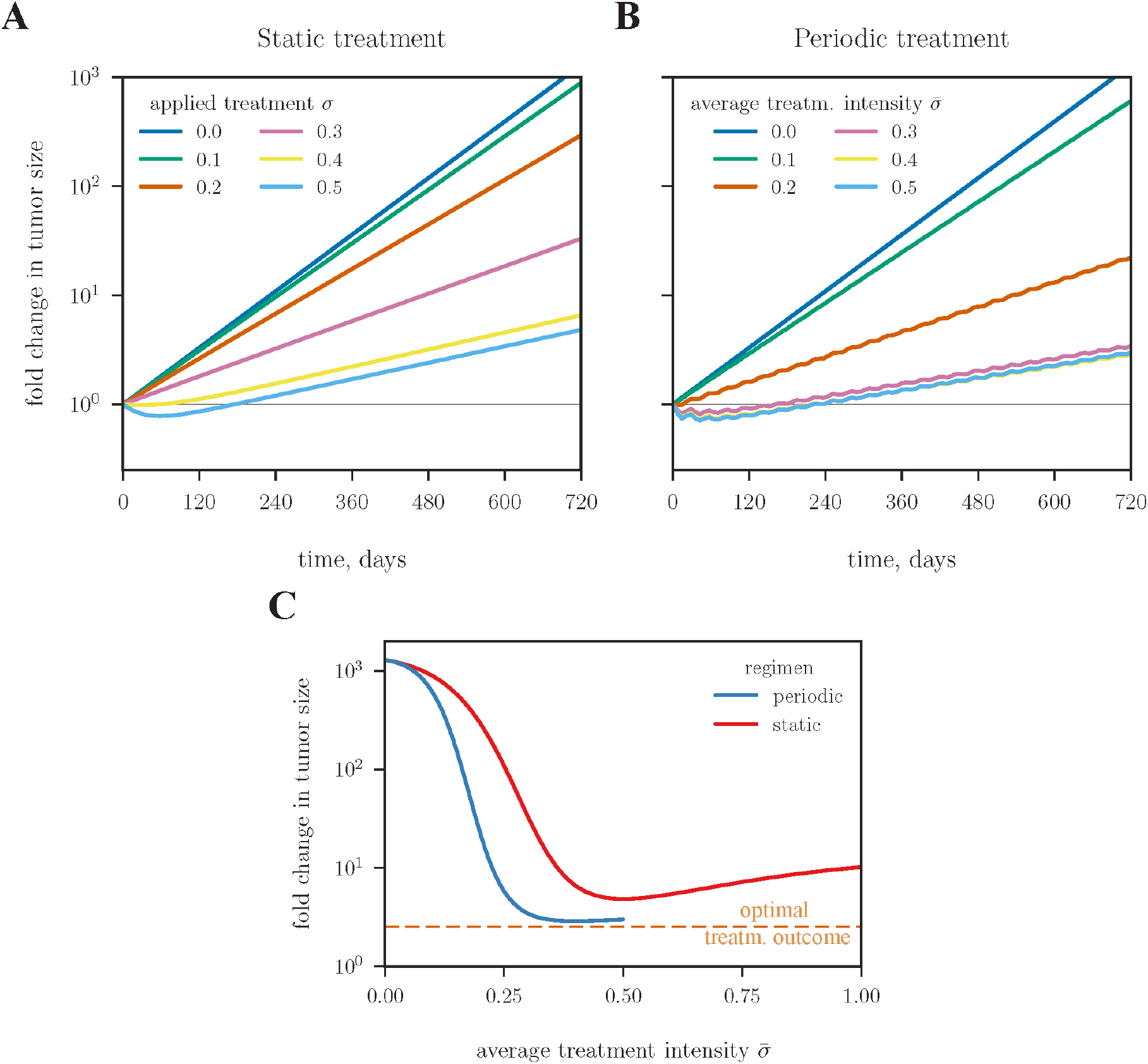
Comparison of static and periodic treatments. (A) Change in tumour size in two years of static treatments. Responses of treatments with six drug intensities are shown in different colours. (B) Tumour dynamics in two years of periodic treatments. Each period consists of 14 days of active phases interleaved with 14 days of drug holidays. To equalize the cumulative drug effect of the two strategies over the entire treatment duration, two treatments are comparable when the drug intensity of the periodic treatment is two-fold as that of the corresponding static treatment. Static treatments with intensities >0.5 (not shown in A) may perform better than periodic regimens due to higher cumulative drug dosage. (C) Change in tumour size after two years of treatment for static (red) and periodic regimen (blue). The dashed orange line indicates the best outcome for the tumour management obtained by solving an optimal control problem.

To compare the outcomes of different treatment regimens, we report the fold change in tumour size after two years. For static treatments with varying intensities (Fig 2C, red curve), the best outcome is reached at an intermediate level of applied treatment intensity (2.3-fold increase at *σ* = 0.52). Treatment intensities higher or lower than the minimizer will lead to larger tumour sizes, yet the level of increase is highly skewed toward left. For example, the treatment of *σ* = 0.1 gives an extremely large final increase in tumour size, while the treatment of a MTD (*σ* = 1.0) leads to a 3.1-fold final increase in tumour size.

Treatment outcomes can also be quantified by the relapse time of regrowing back to its initial size. We confirm again that the optimal setting for static treatments is to apply the intermediate treatment intensity. The maximal relapse time of 6.33 months is achieved at *σ* = 0.59 (Fig S3A). The MTD yields a relapse time of 5.43 months, despite the fact that it reduces the tumour size by the maximal amount of 36% during the initial remission period compared to all other regimens (Fig S3B). The MTD is thus beneficial only in a short-term. The static treatment of low intensity (e.g. *σ* = 0.1) only slows down tumour growth and does not lead to a remission. This confirms that therapy of adequate intensity is required and beneficial. Tumour shrinkage may not predict subsequent outcomes when dynamics of heterogeneous populations are considered.

### 2. Periodic treatment

Tumour relapse is driven by emergence of resistant cells. This process is reversible in our model (Fig S3C), so we may expect improvement in the therapeutic outcomes by leveraging treatment and non-treatment to create a subtle balance of proliferative and resistant cells. The simplest strategy of this kind is a periodic treatment with an equal length of active phase and drug holidays.

Treatments may yield different outcomes simply due to their difference in the cumulative drug quantities administered during the entire episode. To fairly compare treatments with different period lengths and phases (including the static treatments), we normalize drug intensities by the fraction of active phases over the entire episode. For a fixed time horizon *T*, each periodic treatment is characterized by three parameters: the average treatment intensity 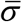, the number of periods of active treatments *K* (*K* = 1,2,…), and the length of each period Δ. The drug intensity administered during the active phase is adjusted to 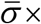 *T* / (total length of active phases). There are two possible scenarios in terms of the phases of the period (Fig 3A). First, the time horizon *T* ends with a drug holiday (*T* − (2*K* − 1)Δ > 0, terminal phase angle *π* ≤ *θ_T_* < 2*π*), then there are *K* full active phases, and the adjusted treatment intensity is given by 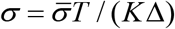. Second, the time horizon *T* ends with an active phase (*T* − (2*K* − 1)Δ < 0, terminal phase angle 0 ≤ *θ_T_* < *π*), then there are (*K* − 1) full drug holidays. The total time of drug administration is *T* − (*K* − 1)Δ, and the adjusted treatment intensity is given by 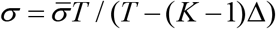. Consequently, the range of all possible adjusted treatment intensities for given *K* is: *σ_K_* ∊ [*σ*_*K*,min_, *σ*_max_], where 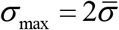 (total period *T* consists of *K* full periods of active phases and drug holidays), and 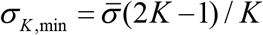 (total period *T* consists of *K* full periods of active phases and (*K* − 1) full periods of drug holidays), see Fig 3B.

**Fig 3:**
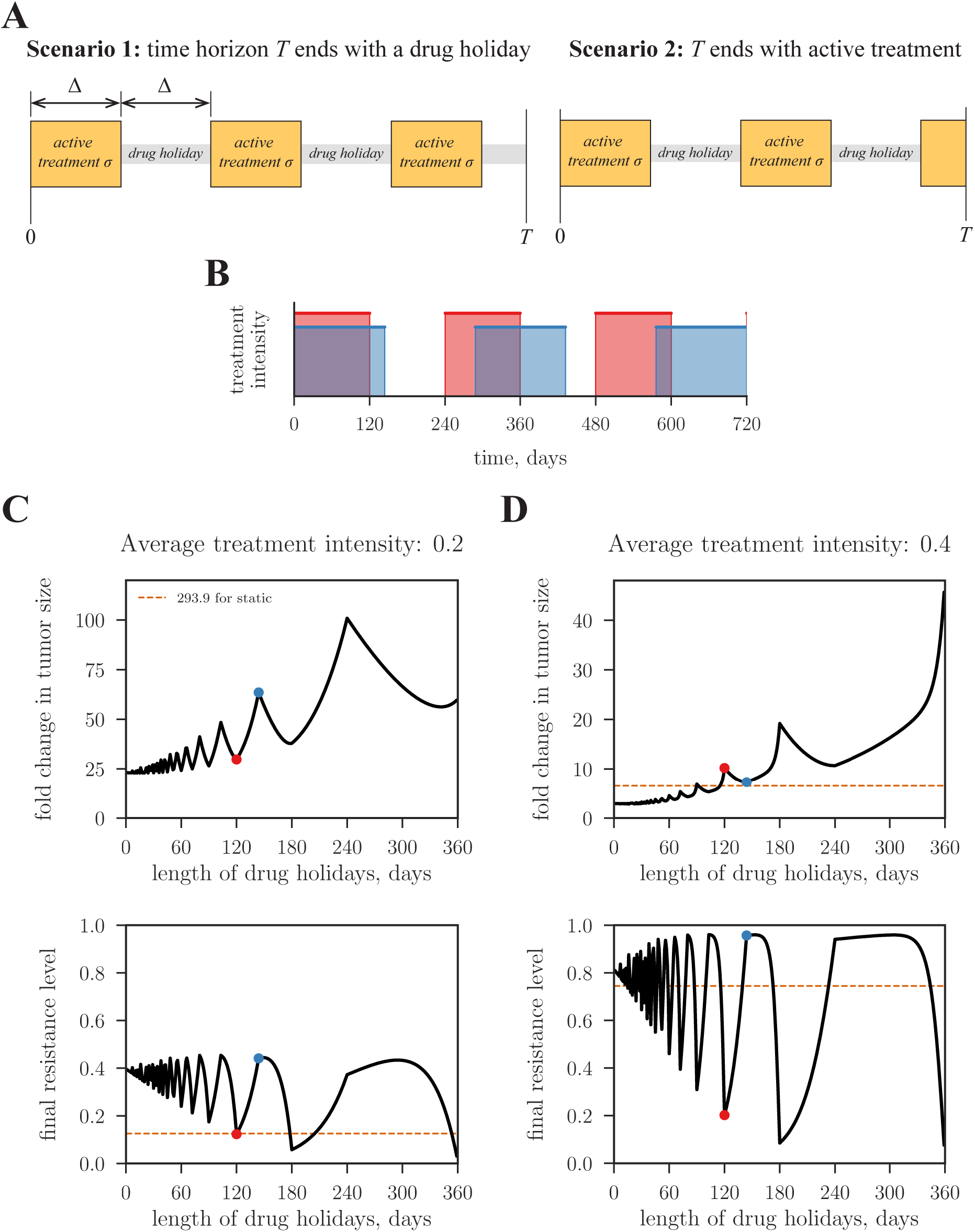
Analysis of periodic treatments. (A) Two scenarios of periodic treatment phases. If the average treatment intensity is constrained by 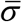, the adjusted treatment intensity administered during the active phase equals 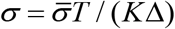 (Scenario 1), and 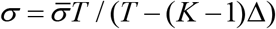 (Scenario 2). *K* = 3 for both scenarios shown in A. (B) Two possible periodic regimens with maximal (red) and minimal (blue) numbers of drug holidays respectively. (C) and (D) shows the outcome of periodic treatments for intermediate and high treatment intensities respectively. Horizontal axes indicate the length of drug holidays per period. Vertical axes indicate the fold change of tumour size after two years. The outcomes of the equivalent static treatments are shown as dashed orange lines. The outcome of the top panel in C is beyond the scale of the *y*-axis (293.9 fold), thus is indicated as a number. Blue and red dots in panels C and D correspond to two periodic schedules shown in panel B.

To assess the influence of treatment schedules on final outcomes, we fix the average dosage intensity and compare tumour size changes in two years with varying period lengths Δ. Dosages of all periodic treatments are adjusted to equalize their cumulative dosages. We first consider treatments of a relatively low intensity 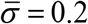 (Fig 3C). The local minima of the tumour size are achieved by applying 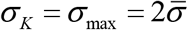 with the terminal phase angle *θ_T_* = 0 (e.g., the red dot in Fig 3C and the red waveform in Fig 3B). In contrast, the schedules that apply 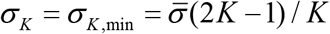 with the terminal phase angle *θ_T_* = *π* (e.g., the blue dot in Fig 3C and the blue waveform in Fig 3B) yield the local maxima in tumour size. All treatment schedules under 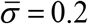 lead to low terminal level of resistance (Fig 3C bottom panel), indicating they are incapable of eliminating the proliferative (sensitive) part of the tumour.

We then consider periodic treatments with a higher average intensity of 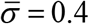 (Fig 3D). In this setting, the outcomes of the two scenarios are the opposite of the previous setting 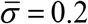. The local minima of the tumour size are achieved by applying 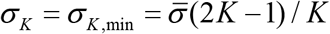 with the terminal phase *θ_T_* = *π* (e.g., the blue dot in Fig 3D and the blue waveform in Fig 3B), whereas the schedules by applying 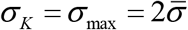 with the terminal phase *θ_T_* = 0 (e.g., the red dot in Fig 3D and the red waveform in Fig 3B) give rise to local maxima in tumour size (e.g., the red dot in Fig 3D and the red waveform in Fig 3B).

The results drawn from these two cases can be summarized as a simple guideline. When the drug appears to be ineffective because only low values of average intensity can be achieved (e.g., 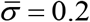), the optimal schedule is to shorten active phases and create an integral number of complete periods, to maximize the average intensity within the active periods. This will amplify the relatively weak intensity within active phases. When the drug appears to be of more effective for pathway inhibition, because sufficiently average intensity is permitted (e.g., 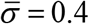), the optimal schedule is to distribute the drug over a longer time span and create a half integral number of complete periods, assuming this is permitted. All the other terminal phases will truncate either an active phase or a drug holiday phase and thus lead to inferior outcomes.

We further find the values of 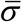 and Δ that jointly optimize the treatment outcome (Fig S4A). Low treatment intensities with 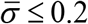 are incapable of controlling tumour growth regardless of treatment schedules. The gradient along Δ is drastically heightened around 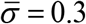. This is exactly when we observe the shift in distribution of local minima and maxima for the fold increase in tumour size (Fig S4C). The global minimum of the tumour size change 2.82 is reached at 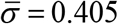 and Δ = 16 days (the red star in Fig S4A). This corresponds to the treatment with 22 active phases and 21 drug holidays during two years of treatment (Fig S4D). However, the terrain of the tumour size change near the global minimum is relatively flat. For instance, the tumour size change is 3.00 when 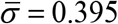 and Δ = 32 days.

Another free parameter of periodic treatments is the duty cycle *a* (length of the active phase of one cycle / length of one cycle). We fix the length of each treatment period to Δ_*c*_ = 2Δ = 32 days and vary *a* and 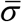 (Fig S5C). Fig S5 indicates the best outcome is achieved at approximately the same treatment as before: *a* = 0.485, 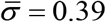, the adjusted treatment intensity during the active phase equals 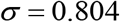, and the fold increase is 2.81. We also obtain similar results when varying the adjusted treatment intensity *σ* rather than the average 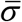 (Fig S6).

### 3. Optimal treatment

Both static and periodic treatments are straightforward to implement but often not optimal in terms of the outcome. Here we define optimality as minimizing the tumour size at a fixed terminal time (or time horizon) *T*. To solve this problem, we apply optimal control theory to update treatment intensities at each moment depending on the tumour state (tumour size and level of resistance). This requires a constant monitoring of the patient, see [16] for discussion. In brief, treatment design is translated into the problem of controlling the temporal function of treatment intensity *σ*(*t*) to minimize the log ratio of final to initial tumour sizes ln(*n*(*T*)/ *n*(0)), subjected to the tumour population dynamics (equation 1). Solution of the optimal control problem is described in Appendix B and based on the method of generalized characteristics [49–51].

The optimal *σ*(*t*) is determined by both the initial proportion of resistant cells and the length of the time horizon. Fig 4 illustrates optimal trajectories of three initial conditions. (*i*) The time horizon *T* is shorter than a threshold: *T* = *T*_−_ < *T*_0_, and initially the proportion of resistant cells is 0. The optimal treatment applies a dosage close to MTD for the whole period. Proportion of resistance cells increases over time and reaches a level of about 75% for given baseline parameters at the terminal point. (*ii*) *T* is longer than the same threshold: *T* = *T*_+_ > *T*_0_, and initially the proportion of resistant cells is 0. The optimal treatment comprises three stages. It starts with a high intensity close MTD for about one month, then sharply lowers the dose to a moderate level till about one month before the terminal point, and finally resumes the high dosage till the end. Proportion of resistance cells climbs up and reaches about 75% in the first stage, maintains at this level in the second stage, and further increases again in the third stage. (*iii*) *T* is longer than the same threshold: *T* = *T*_+_ > *T*_0_, and initially the proportion of resistant cells is 1. The optimal treatment also consists of three stages. In the first stage the drug is not administered. Proportion of resistant cells thus decreases to 75%. Treatment intensities and resistance trajectories in the second and third stages coincide with case (*ii*). Importantly, a dose-sparing regimen in the middle of (*ii*) and (*iii*) concurs with the periodic treatment in its efficiency by keeping the balance between sensitive and resistant parts of the tumour. This prepares the patient for the final stage of the treatment when the sensitive part of the tumour is eradicated with greater efficiency. Overall, optimal treatments aim for establishing and maintaining a fixed balance between proliferative and resistant cells as long as possible until near the terminal point, and then switch to the maximal dosage throughout the remaining time to eradicate as many proliferative cells as possible (curves *ii* and *iii* in Fig 4; Fig 5B). Yet when the time horizon is short, the long term benefit of a balanced population is no longer relevant, and the optimal treatment is to reduce the current tumour size by administering the maximal dosage (curve *i* in Fig 4; Fig 5B). The clinically relevant time horizon may depend on other factors such as the emergence of genetically distinct subclones with different properties.

**Fig 4:**
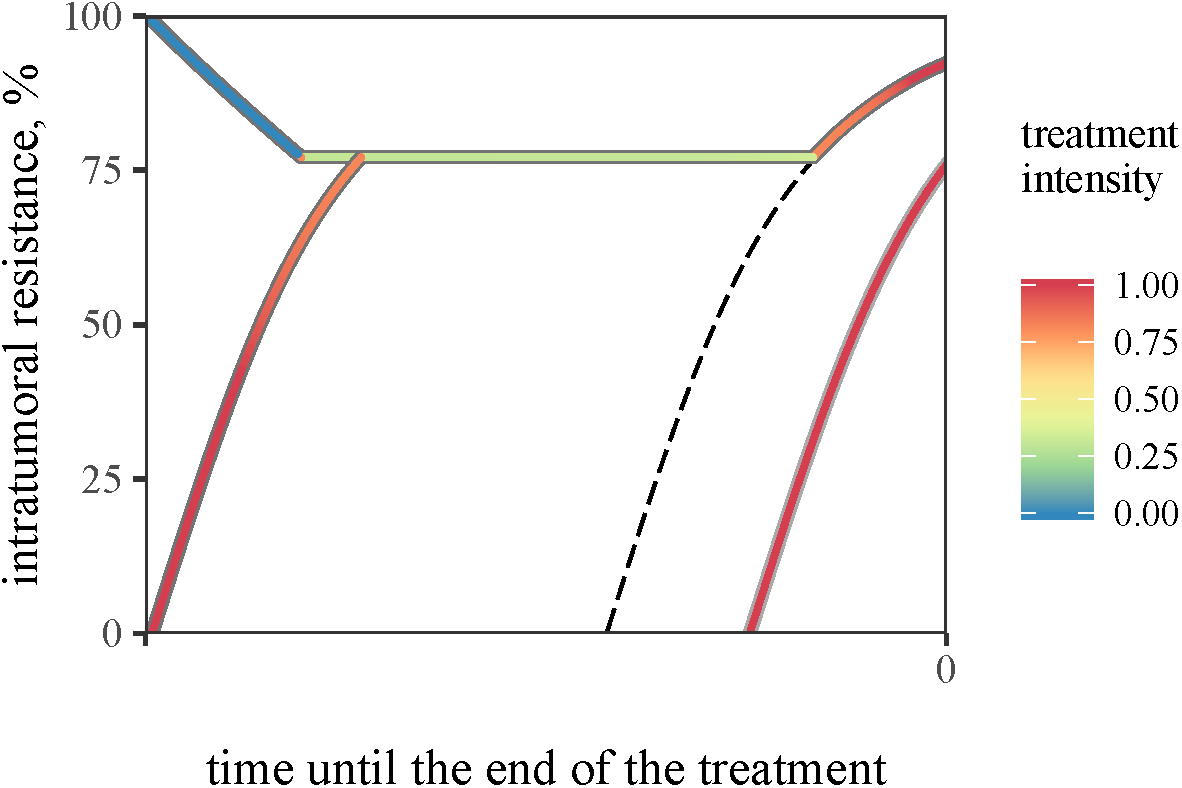
Optimal treatment intensity is determined by the current level of resistance and the total time span of treatment. The temporal axis marks the direction from the start of a treatment (a positive number) to the terminal point of treatment. The threshold value *T*_0_ = 1.75 months separates the trajectories with and without a dose-sparing regimen whose trajectory is marked by the dashed line. Trajectories of three regimens are illustrated (see description in the text). Trajectory colours indicate the applied treatment intensity (legend).

**Fig 5:**
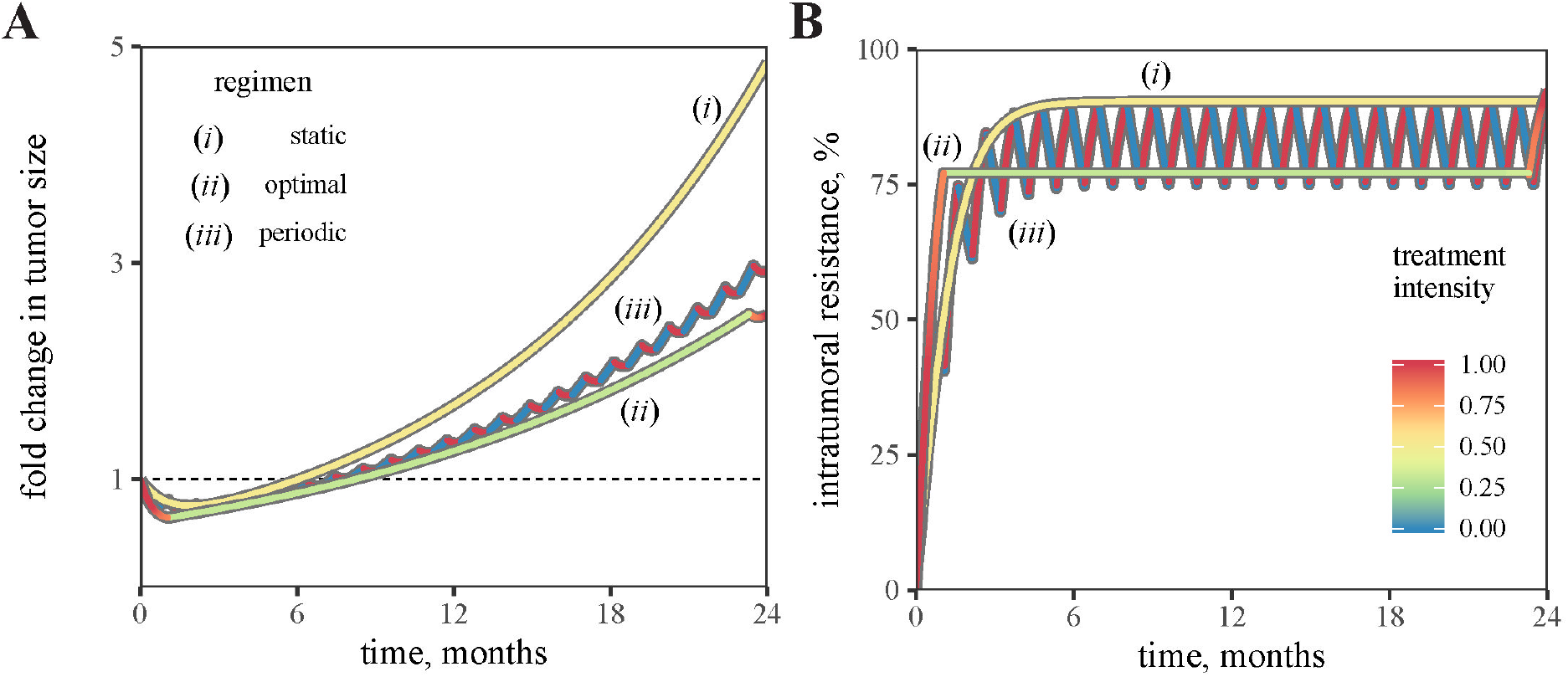
Comparison of three different treatment schedules in terms of fold-change in tumour size (A) and dynamics of intratumoral resistance (B). The dashed horizon indicates the fold-change equal one. Line colours indicate the applied treatment intensity.

### 4. Comparison of different treatments

The performances of the three aforementioned treatment strategies conform with the following order: best static treatment ≤ best periodic treatment ≤ optimal treatment. Superiority of periodic over static treatments is illustrated in Fig 2. The treatment derived from the optimal control theory is superior to all dynamic treatments including periodic treatments. To quantitatively compare their performances, we fix the time horizon to two years, set the best static treatment intensity to *σ* = 0.52, the best periodic treatment intensity to *σ* = 0.8 and period to 16 days according to Fig S4C, and find the optimal dynamic treatment by solving the optimal control problem. Fig 5 shows the comparison outcomes of those three treatments. The terminal tumour sizes (relative the initial tumour size) are consistent with the aforementioned order. The static treatment yields a poor outcome (4.85-fold increase of tumour size after two years). The periodic treatment gives a much better result (2.93-fold increase after two years), which is just marginally inferior to the minimally achievable estimate obtained from the optimal treatment (2.52-fold change after two years). Notice that the optimal strategy keeps the tumour size higher than in the periodic treatment until about one month before the terminal time. This counter-intuitive action provides a balance between the sensitive and resistant cells and allows more efficient reduction of the tumour size at the final stage. The population composition trajectories of the three treatments are shown in Fig 5B. Proportion of resistant cells steadily increases to a fixed value and maintains onward in the static treatment due to the constant administration of the drug. Proportion of resistant cells in the periodic treatment undergoes an initial transient stage and oscillates around a fixed value, synchronous with the period of the treatment. Proportion of resistant cells in the optimal treatment sharply reaches a fixed level, remains invariant most of the time, and suddenly increases in the last stage. This pattern follows exactly the scenarios described in Fig 4.

### 5. Sensitivity analysis

Eleven baseline parameter values in the model (Table 1) are not guaranteed to be accurate and unique. To investigate the influence of parameter values in optimal treatment outcomes, we assess the fold change in tumour size after two years by varying parameter values. Fig 6A shows the effect of variation in characteristic switching times between the main and alternative pathways, when the cost of resistance *c* = 4%. The tumour does not shrink if the inverse switch from the alternative to the main pathway is slower than the direct switch from the main to the alternative pathway (the red-yellow region above the solid black line), while remission can be achieved if the reciprocal relation between the two switching times holds (the blue region below the solid black line). However, the result depends on the cost of resistance: a higher cost induces slower proliferation of resistant cells and thus accommodates a wider range of switching times leading to tumour reduction (Fig 6C), while a lower cost has the opposite effect (Fig 6B). We further investigate how the optimal proportion of resistant and sensitive cells depends on aforementioned parameters (Fig S7). The optimal proportion of resistant cells is positively correlated with 1/ *μ* (Fig S7B) and negatively correlated with 1/ *μ* (Fig S7A). We also notice that the optimal proportion is below 50% only when *μ* is four times slower than 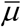 (area below dashed line in Fig S7C).

**Fig 6:**
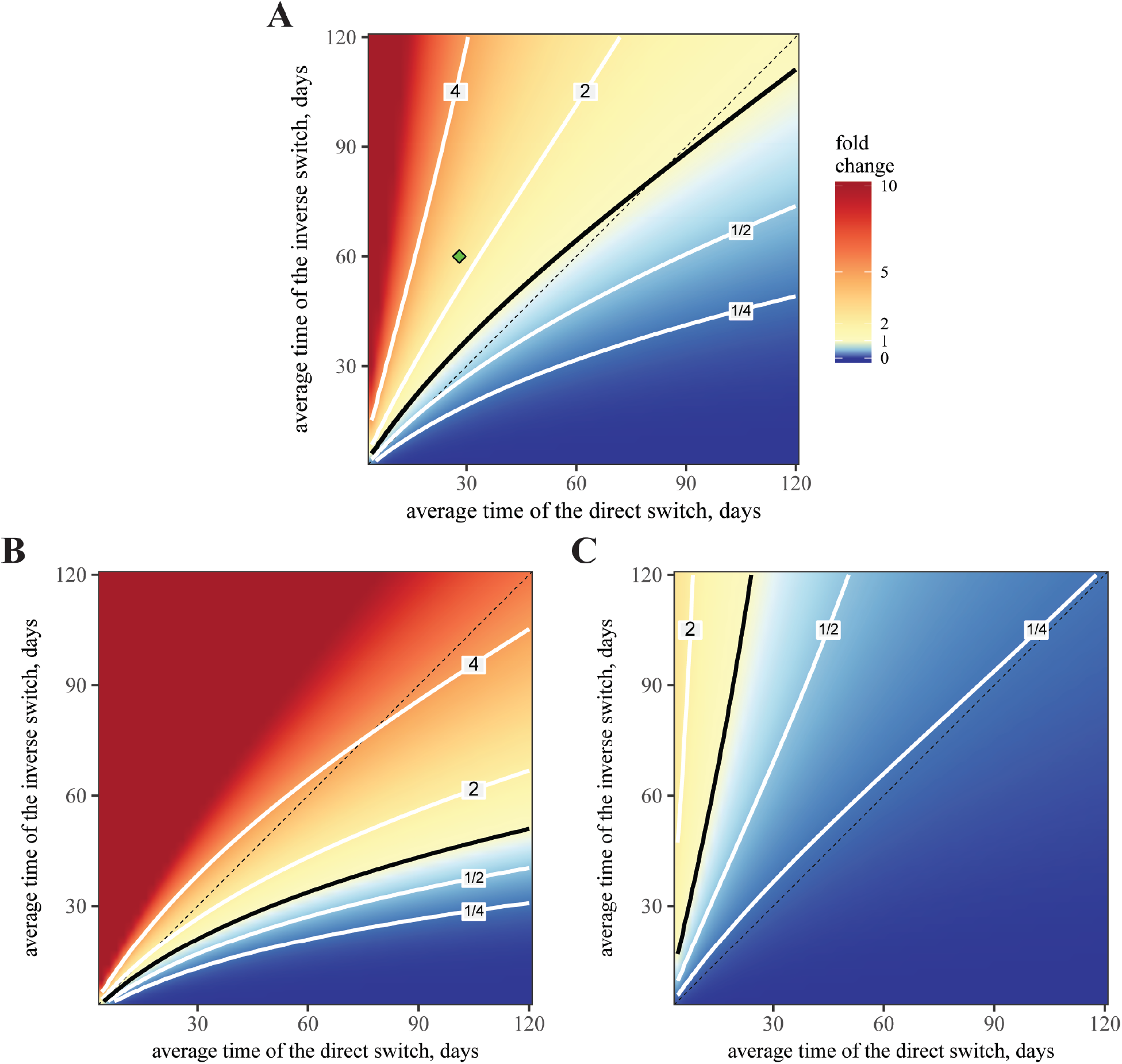
Sensitivity of tumour size after two years of optimally designed treatment. The varied parameters are characteristic switching times *μ*^−1^ and 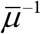 (horizontal and vertical axis respectively). Three panels refer to different values of resistance cost: *c* = 4% (baseline, A), 2% (B), 6% (C). Other parameters are fixed according to Table 1. The contour line for the fold change equal to one is indicated by solid black, four other contours are shown in grey and correspond to the fold change in the panel to each line. Diagonal for 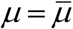 is shown in dashed black. The location of the baseline parameters is indicated by the green point in A.

Fig S8 shows the variation of tumour size with respect to other model parameters. The most sensitive parameters are the cost of resistance *c* (Fig S8A) and the robustness parameter *κ* (Fig S8E). The latter indicates how likely the main pathway switches between its on- and off-states due to internal stochastic effects. The system possessing the baseline *κ* value used in our simulation yields near the smallest tumour size (green dot in Fig S8E). A more unstable main pathway (smaller *κ* values) impairs the effect of drug holidays as a sensitive cell will randomly drift to the resistant state with high probability. Reciprocally, a more stable main pathway (higher *κ* values) deteriorates the effect of treatment as a cell is not responsive to the pathway inhibitor. Thus time-varying treatments are effective only in a narrow range of pathway robustness.

## Discussion

Both optimal and periodic treatments outperform the static treatment by more than two folds in terms of the tumour size after two years (2.52-fold, 2.93-fold and 4.85-fold respectively). Efficacy of periodic treatments was discussed in prior studies [16,52]. Drug addiction is one possible cause [33]: resistant subclones not only tolerate the administered drug but also depend on it. Drug holidays in these cases deplete the “nutrient” supply and reduce the resistant subclone population. Gatenby *et al*. [53] considered a more general “adaptive therapy” as means to maintain proper balance of sensitive and resistant subclone populations undergoing competition. Our simulation outcomes corroborate the superiority of periodic treatments and concur with the prior discussions about their benefits, albeit the proposed mechanisms causing the benefits are different. Those mechanisms may co-exist and can be all tackled by periodic treatments, at least in the setting of a single therapy as modelled here. In spite of a good approximation to the global optimum and simplicity of implementation, the best periodic treatment is still marginally inferior to the optimal treatment strategy. In principle, one should always adopt the treatment strategy that yields the best outcome. In practice, physicians have to consider multiple factors when deciding the treatment including cost-effectiveness, the higher risk of medical errors when implementing complex recommendations, and the feasibility of repeated tumour sampling or liquid biopsy to provide accurate input data to correctly design and execute an optimal control algorithm. Both periodic and optimal treatments likely require substantial modifications. Traditionally in clinical studies the current agent is abandoned when there is a 20% increase in the sum of the longest linear dimension of large measurable lesions, corresponding to a 73% increase in volume [54]. Such a practice does not allow for periodic treatments. Also, the optimal treatment in our model requires continuous monitoring of the tumour population composition. If such monitoring is either costly or infeasible, then a properly designed periodic treatment is a reasonable surrogate for the optimal treatment. Dynamic treatments of cancer require biomarkers for functional states that can be continuously and non-invasively monitored. This may be a greater challenge for complex gene expression changes and their associated physiological changes than it is for mutations for which highly sensitive polymerase chain reaction techniques are available.

Intriguingly, the best treatment plans of all three strategies attempt to establish and maintain an optimal proportion of sensitive and resistant cells (fraction of sensitive cells is about 75% in our simulations) throughout the entire period (Fig 5B). The best static treatment drives the population toward a value above such optimal composition. The best periodic treatment quickly leads the population to the optimal composition and then makes it oscillate around this value. The global optimal treatment quickly forces the population toward the optimal composition, alters the dosage to maintain it, and finally maximizes the dosage for final most efficient curbing of the tumour at the very end. The outcome of a treatment strategy seems to depend critically on its controllability to reach this composition as quick as possible and stay there as long as possible. In this sense, the static treatment is a poor controller, because it cannot reach that target. Both periodic and optimal treatments are good controllers as they quickly reach the target value and maintain it onward. Curiously, while the total population under the best periodic treatment steadily increases with small ripples, the total population under the optimal treatment grows rapidly and finally plunges to a much lower value (Fig 5A). This is because the higher growth rate allowed by the optimal treatment leads to a more optimal balance of sensitive and resistant cells for efficient final curbing of the tumour. This also identifies a possible hurdle in implementing an optimal treatment regimen into clinical practice because its performance at the initial state is even worse than with a static treatment, and this is likely to result in earlier termination of the therapy per current paradigms (Fig 5A). However, design of optimal treatment may help to estimate performance of other suboptimal treatments such as periodic or adaptive regimens. Thus, the solution of optimal control problems for tumour management may contribute valuable information even if not applied in practice.

Many of the parameter estimates such as the characteristic time of switching between two genetic programs are subjected to uncertainty. Despite impressive progress in the field, recent studies [8,14] indicated that the technology of single-cell transcriptomics still does not allow rigorous measurement of the kinetics of a change in growth rates of sensitive and resistant cells due to epigenetic reprogramming. Laboratory studies in cell lines may not reflect observations of net growth rates of tumours in patients (*cf* reported proliferation rates in [13] and [55]). This causes difficulties for model inference. Based on this, we first proposed an estimate of characteristic switching times and then implemented a sensitive analysis to verify the robustness of our outcomes. We anticipate in the future that laboratory model systems will be developed that enable testing and refinement of our findings.

We assume that reprogrammable cell states require not only an instant change in activity of both pathways (with faster time scales), but also its “hardwiring” in the physiology of a cell [25]. For example, a recent study [8] implements a special state called a meta-resistant state that still does not guarantee a permanent resistant state, but remains reversible, similar to our model formulation (*cf* Fig 1B and C).

The model in this study captures aspects of the reversible process of drug resistance. By integrating with our prior work for the irreversible process of drug resistance [30,31], we plan to build a relatively complete model capturing both reversible and irreversible processes and design the treatment strategies accordingly. Outgrowth of rare subclonal resistance mutations or acquisition of new resistance mutation may occur on a longer timescale than the phenomena discussed herein. Extension in multiple directions is required, such as incorporating the activities of the major cancer pathways in the cellular internal states, expanding the drugs and treatment options, and including molecular-level resistance mechanisms. However, such extension will also substantially increase the model complexity and data requirements, a particular problem for clinical translation. A principled method to balance the required features of the model and their associated data requirements, as well as specific methods for dealing with uncertainty and incomplete information, remain critical tasks.

## Acknowledgments

Code for all calculations, and for producing all of the figures, is available at http://tiny.cc/AkhmKim18Scripts. ARA performed computer simulations using the facilities of the SHARCNET (www.sharcnet.ca) and Compute/Calcul Canada. ARA’s postdoctoral stay was financially supported by Academia Sinica (Taiwan). PT and JWK were supported by the United States NIH/NCI grant U01CA217885. CHY was supported by Career Development Award 104-CDA-M04 from Academia Sinica and Ministry of Science and Technology (MOST) of Taiwan grant 103-2118-M-001-011-MY2.

## References

1. Garraway LA, Jänne PA. 2012 Circumventing cancer drug resistance in the era of personalized medicine. Cancer Discovery 2, 214–26. (doi:10.1158/2159-8290.CD-12-0012)

2. Bozic, Nowak MA. 2017 Resisting Resistance. Annu. Rev. Cancer Biol. 1, 203–221. (doi:10.1146/annurev-cancerbio-042716-094839)

3. Hu X, Zhang Z. 2016 Understanding the Genetic Mechanisms of Cancer Drug Resistance Using Genomic Approaches. Trends Genet. 32, 127–137. (doi:10.1016/j.tig.2015.11.003)

4. Sharma P, Hu-Lieskovan S, Wargo JA, Ribas A. 2017 Primary, Adaptive, and Acquired Resistance to Cancer Immunotherapy. Cell 168, 707–723. (doi:10.1016/j.cell.2017.01.017)

5. Dagogo-Jack I, Shaw AT. 2017 Tumour heterogeneity and resistance to cancer therapies. Nat. Rev. Clin. Oncol. 15, 81–94. (doi:10.1038/nrclinonc.2017.166)

6. Greaves M, Maley CC. 2012 Clonal evolution in cancer. Nature 481, 306–313. (doi:10.1038/nature10762)

7. Williams MJ, Werner B, Heide T, Curtis C, Barnes CP, Sottoriva A, Graham TA. 2018 Quantification of subclonal selection in cancer from bulk sequencing data. Nat. Genet. 50, 895–903. (doi:10.1038/s41588-018-0128-6)

8. Shaffer SM et al. 2017 Rare cell variability and drug-induced reprogramming as a mode of cancer drug resistance. Nature 546, 431–435. (doi:10.1038/nature22794)

9. Stites EC. 2012 The Response of Cancers to BRAF Inhibition Underscores the Importance of Cancer Systems Biology. Sci. Signal. 5, pe46–pe46. (doi:10.1126/scisignal.2003354)

10. Gerlinger M et al. 2012 Intratumor Heterogeneity and Branched Evolution Revealed by Multiregion Sequencing. N. Engl. J. Med. 366, 883–892. (doi:10.1056/NEJMoa1113205)

11. Hangauer MJ et al. 2017 Drug-tolerant persister cancer cells are vulnerable to GPX4 inhibition. Nature (doi:10.1038/nature24297)

12. Sharma SV et al. 2010 A Chromatin-Mediated Reversible Drug-Tolerant State in Cancer Cell Subpopulations. Cell 141, 69–80. (doi:10.1016/j.cell.2010.02.027)

13. Sun C et al. 2014 Reversible and adaptive resistance to BRAF(V600E) inhibition in melanoma. Nature 508, 118–122. (doi:10.1038/nature13121)

14. Tirosh I et al. 2016 Dissecting the multicellular ecosystem of metastatic melanoma by single-cell RNA-seq. Science 352, 189–196. (doi:10.1126/science.aad0501)

15. Kuczynski EA, Sargent DJ, Grothey A, Kerbel RS. 2013 Drug rechallenge and treatment beyond progression—implications for drug resistance. Nat. Rev. Clin. Oncol. 10, 571–587. (doi:10.1038/nrclinonc.2013.158)

16. Fischer A, Vázquez-García I, Mustonen V. 2015 The value of monitoring to control evolving populations. Proc. Natl. Acad. Sci. 112, 1007–1012. (doi:10.1073/pnas.1409403112)

17. Beerenwinkel N, Schwarz RF, Gerstung M, Markowetz F. 2015 Cancer Evolution: Mathematical Models and Computational Inference. Syst. Biol. 64, e1–e25. (doi:10.1093/sysbio/syu081)

18. Michor F, Beal K. 2015 Improving Cancer Treatment via Mathematical Modeling: Surmounting the Challenges Is Worth the Effort. Cell 163, 1059–1063. (doi:10.1016/j.cell.2015.11.002)

19. Ashcroft P, Michor F, Galla T. 2015 Stochastic Tunneling and Metastable States During the Somatic Evolution of Cancer. Genetics 199, 1213–1228. (doi:10.1534/genetics.114.171553)

20. Sprouffske K, Pepper JW, Maley CC. 2011 Accurate Reconstruction of the Temporal Order of Mutations in Neoplastic Progression. Cancer Prev. Res. (Phila. Pa.) 4, 1135–1144. (doi:10.1158/1940-6207.CAPR-10-0374)

21. Dingli D, Chalub FACC, Santos FC, Van Segbroeck S, Pacheco JM. 2009 Cancer phenotype as the outcome of an evolutionary game between normal and malignant cells. Br. J. Cancer 101, 1130–1136. (doi:10.1038/sj.bjc.6605288)

22. You L, Brown JS, Thuijsman F, Cunningham JJ, Gatenby RA, Zhang J, Staňková K. 2017 Spatial vs. non-spatial eco-evolutionary dynamics in a tumor growth model. J. Theor. Biol. 435, 78–97. (doi:10.1016/j.jtbi.2017.08.022)

23. Klement GL. 2016 Eco-evolution of cancer resistance. Sci. Transl. Med. 8, 327fs5–327fs5. (doi:10.1126/scitranslmed.aaf3802)

24. Taylor-King JP, Baratchart E, Dhawan A, Coker EA, Rye IH, Russnes H, Chapman SJ, Basanta D, Marusyk A. 2018 Simulated ablation for detection of cells impacting paracrine signalling in histology analysis. Math. Med. Biol. J. IMA (doi:10.1093/imammb/dqx022)

25. Bacevic K et al. 2017 Spatial competition constrains resistance to targeted cancer therapy. Nat. Commun. 8. (doi:10.1038/s41467-017-01516-1)

26. Kim E, Kim J-Y, Smith MA, Haura EB, Anderson ARA. 2018 Cell signaling heterogeneity is modulated by both cell-intrinsic and -extrinsic mechanisms: An integrated approach to understanding targeted therapy. PLOS Biol. 16, e2002930. (doi:10.1371/journal.pbio.2002930)

27. Marusyk A, Almendro V, Polyak K. 2012 Intra-tumour heterogeneity: a looking glass for cancer? Nat. Rev. Cancer 12, 323–334. (doi:10.1038/nrc3261)

28. Kolch W, Halasz M, Granovskaya M, Kholodenko BN. 2015 The dynamic control of signal transduction networks in cancer cells. Nat. Rev. Cancer 15, 515–527. (doi:10.1038/nrc3983)

29. Jolly MK, Kulkarni P, Weninger K, Orban J, Levine H. 2018 Phenotypic Plasticity, Bet-Hedging, and Androgen Independence in Prostate Cancer: Role of Non-Genetic Heterogeneity. Front. Oncol. 8. (doi:10.3389/fonc.2018.00050)

30. Beckman RA, Schemmann GS, Yeang C-H. 2012 Impact of genetic dynamics and singlecell heterogeneity on development of nonstandard personalized medicine strategies for cancer. Proc. Natl. Acad. Sci. 109, 14586–14591. (doi:10.1073/pnas.1203559109)

31. Yeang C-H, Beckman RA. 2016 Long range personalized cancer treatment strategies incorporating evolutionary dynamics. Biol. Direct 11. (doi:10.1186/s13062-016-0153-2)

32. Sosman JA et al. 2012 Survival in BRAF V600–Mutant Advanced Melanoma Treated with Vemurafenib. N. Engl. J. Med. 366, 707–714. (doi:10.1056/NEJMoa1112302)

33. Das Thakur M, Salangsang F, Landman AS, Sellers WR, Pryer NK, Levesque MP, Dummer R, McMahon M, Stuart DD. 2013 Modelling vemurafenib resistance in melanoma reveals a strategy to forestall drug resistance. Nature 494, 251–255. (doi:10.1038/nature11814)

34. Siegel RL, Miller KD, Jemal A. 2018 Cancer statistics, 2018: Cancer Statistics, 2018. CA. Cancer J. Clin. 68, 7–30. (doi:10.3322/caac.21442)

35. Akbani R et al. 2015 Genomic Classification of Cutaneous Melanoma. Cell 161, 1681–1696. (doi:10.1016/j.cell.2015.05.044)

36. Dummer R et al. 2018 Encorafenib plus binimetinib versus vemurafenib or encorafenib in patients with BRAF-mutant melanoma (COLUMBUS): a multicentre, open-label, randomised phase 3 trial. Lancet Oncol. 19, 603–615. (doi:10.1016/S1470-2045(18)30142-6)

37. Larkin J et al. 2014 Combined vemurafenib and cobimetinib in BRAF-mutated melanoma. N. Engl. J. Med. 371, 1867–1876. (doi:10.1056/NEJMoa1408868)

38. Johannessen CM et al. 2010 COT drives resistance to RAF inhibition through MAP kinase pathway reactivation. Nature 468, 968–972. (doi:10.1038/nature09627)

39. Montagut C et al. 2008 Elevated CRAF as a Potential Mechanism of Acquired Resistance to BRAF Inhibition in Melanoma. Cancer Res. 68, 4853–4861. (doi:10.1158/0008-5472.CAN-07-6787)

40. Nazarian R et al. 2010 Melanomas acquire resistance to B-RAF(V600E) inhibition by RTK or N-RAS upregulation. Nature 468, 973–977. (doi:10.1038/nature09626)

41. Poulikakos PI et al. 2011 RAF inhibitor resistance is mediated by dimerization of aberrantly spliced BRAF(V600E). Nature 480, 387–390. (doi:10.1038/nature10662)

42. Konieczkowski DJ et al. 2014 A Melanoma Cell State Distinction Influences Sensitivity to MAPK Pathway Inhibitors. Cancer Discov. 4, 816–827. (doi:10.1158/2159-8290.CD-13-0424)

43. Müller J et al. 2014 Low MITF/AXL ratio predicts early resistance to multiple targeted drugs in melanoma. Nat. Commun. 5, 5712. (doi:10.1038/ncomms6712)

44. Kemper K, de Goeje PL, Peeper DS, van Amerongen R. 2014 Phenotype Switching: Tumor Cell Plasticity as a Resistance Mechanism and Target for Therapy. Cancer Res. 74, 5937–5941. (doi:10.1158/0008-5472.CAN-14-1174)

45. Hanahan D, Weinberg RA. 2000 The Hallmarks of Cancer. Cell 100, 57–70. (doi:10.1016/S0092-8674(00)81683-9)

46. Krapivsky P, Redner S, Ben-Naim E. 2010 A kinetic view of Statistical Physics. Cambridge: Cambridge University Press.

47. Chmielecki J et al. 2011 Optimization of Dosing for EGFR-Mutant Non-Small Cell Lung Cancer with Evolutionary Cancer Modeling. Sci. Transl. Med. 3, 90ra59–90ra59. (doi:10.1126/scitranslmed.3002356)

48. Nowak MA. 2006 Evolutionary dynamics: exploring the equations of life. Cambridge, Mass: Belknap Press of Harvard University Press.

49. Melikyan A. 1998 Generalized Characteristics of First Order PDEs. Boston, MA: Birkhäuser Boston. (doi:10.1007/978-1-4612-1758-9)

50. Melikyan AA, Ovseevich AI. 1984 Hamiltonian systems with a specified invariant manifold and some of their applications. J. Appl. Math. Mech. 48, 140–145. (doi:10.1016/0021-8928(84)90079-0)

51. Melikyan AA, Ovseevich AI. 2011 Universal surfaces and smooth solutions of Bellman’s equations. Russ. J. Math. Phys. 18, 176–182. (doi:10.1134/S1061920811020063)

52. Foo J, Michor F. 2009 Evolution of Resistance to Targeted Anti-Cancer Therapies during Continuous and Pulsed Administration Strategies. PLoS Comput. Biol. 5, e1000557. (doi:10.1371/journal.pcbi.1000557)

53. Gatenby RA, Silva AS, Gillies RJ, Frieden BR. 2009 Adaptive Therapy. Cancer Res. 69, 4894–4903. (doi:10.1158/0008-5472.CAN-08-3658)

54. Eisenhauer EA et al. 2009 New response evaluation criteria in solid tumours: Revised RECIST guideline (version 1.1). Eur. J. Cancer 45, 228–247. (doi:10.1016/j.ejca.2008.10.026)

55. Bozic I et al. 2013 Evolutionary dynamics of cancer in response to targeted combination therapy. eLife 2. (doi:10.7554/eLife.00747)

56. Bozic I et al. 2010 Accumulation of driver and passenger mutations during tumor progression. Proc. Natl. Acad. Sci. 107, 18545–18550. (doi:10.1073/pnas.1010978107)

57. Miyamoto T, Furusawa C, Kaneko K. 2015 Pluripotency, Differentiation, and Reprogramming: A Gene Expression Dynamics Model with Epigenetic Feedback Regulation. PLOS Comput. Biol. 11, e1004476. (doi:10.1371/journal.pcbi.1004476)

58. Iwasa Y, Nowak MA, Michor F. 2006 Evolution of Resistance During Clonal Expansion. Genetics 172, 2557–2566. (doi:10.1534/genetics.105.049791)

