## Supporting Materials for "Modelling bistable tumour population dynamics to design effective treatment strategies"

### Modelling bistable tumour population dynamics and designing their effective treatments

#### Supplementary figures

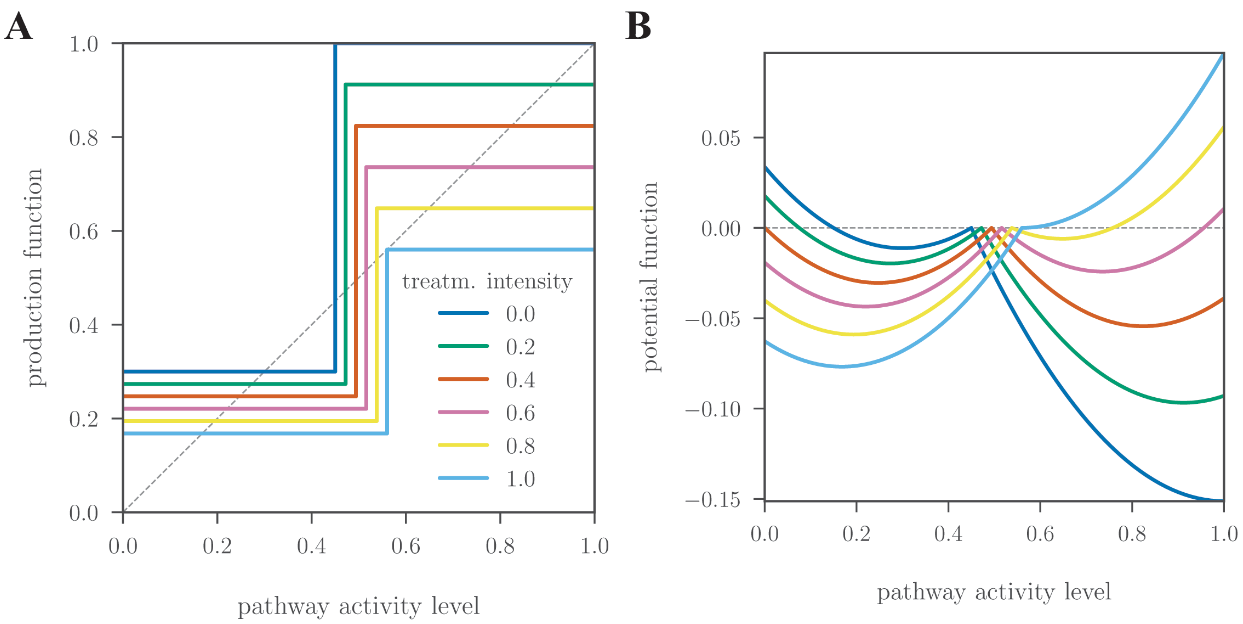


**Fig S1: Production function (A) and corresponding set of potentials (B) under varied treatment intensities.** The grey dashed in B indicates the zero level in the potential.

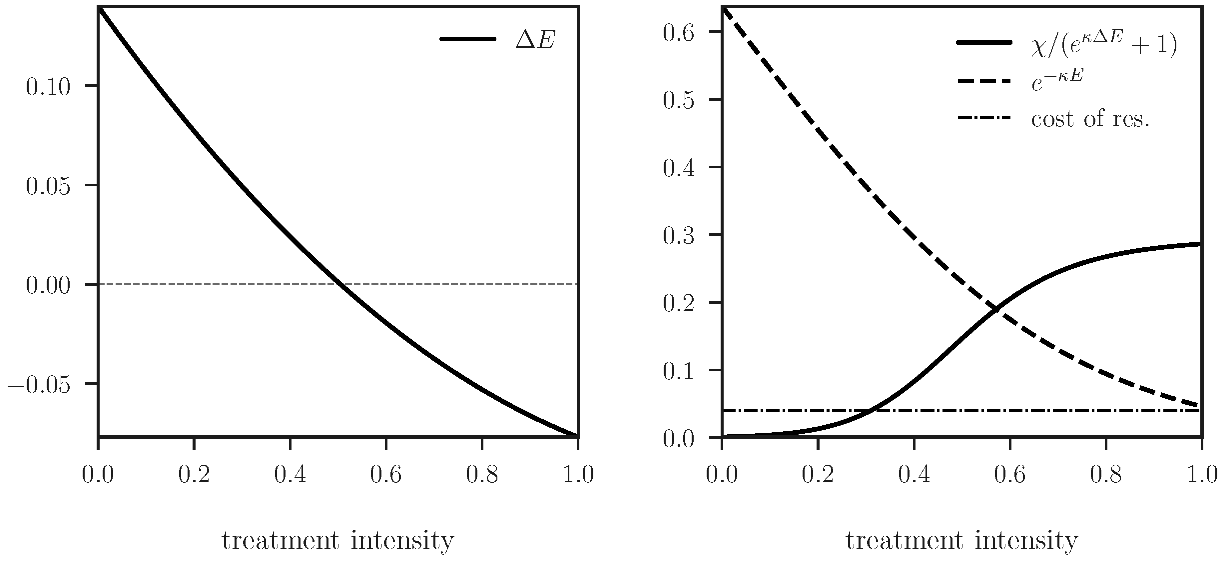


**Fig S2: Main characteristics of the equations** Error! Reference source not found.**–(11) as the functions of treatment intensity.**

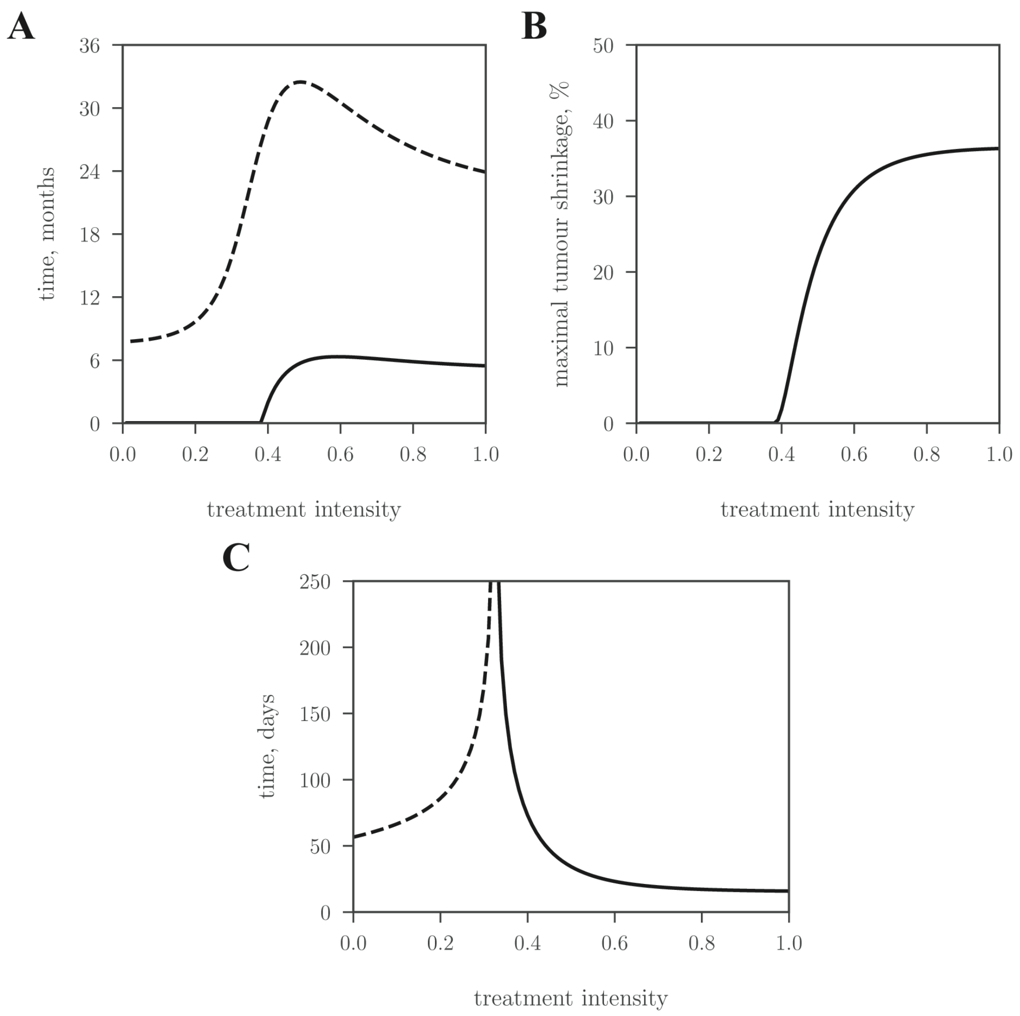


**Fig S3: (A)** Relapse times of returning to the initial size for tumours under static treatments (the solid line). For comparison, the dashed line indicates the time necessary for a 10-fold increase in size. **(B)** Maximal tumour shrinkage observed during the static treatment. **(C)** Time necessary to acquire 50%-level of resistance starting from the initial 1% (solid) vs. the time necessary to lower the resistance level from the initial 99% to 50% (dashed) under the static treatment regimen.

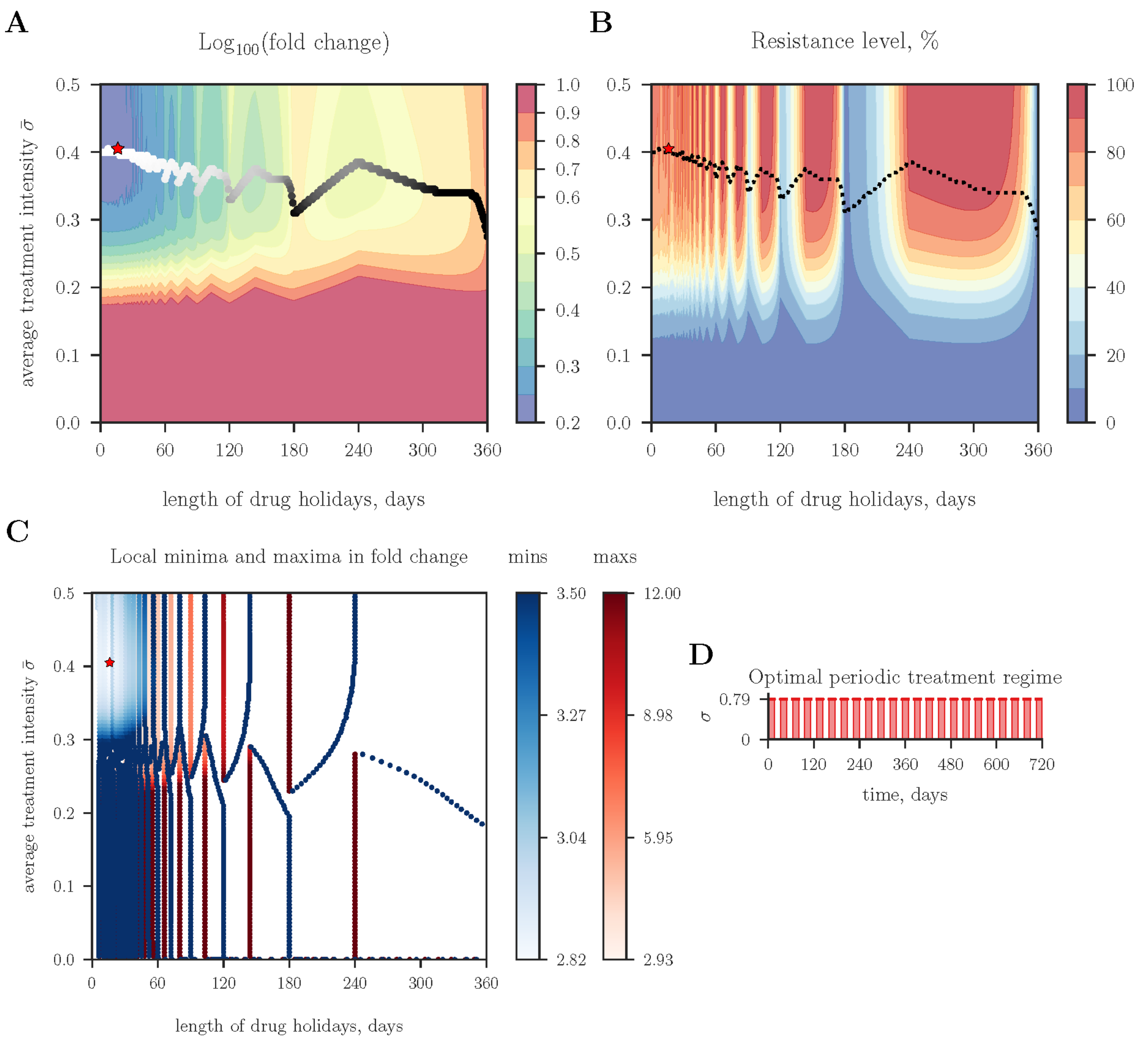


**Fig S4: Analysis of periodic treatment regimen with equal durations of active phases and drug holidays.** All treatments are compared in terms of the average treatment intensity (or cumulative treatment intensity), see *y*-axis. The fold change in tumour size after two years of treatment (A), and the final resistance level within a tumour (B) are characterized by repeating patterns. A line of dots with gradient grey represents the minima in fold change for fixed length of active phases/drug holidays and varied average treatment intensity. The red star indicates the global minimum and relates to the schedule shown (D). (C) shows the change of local minima (gradient of blue) and maxima (gradient of red) in fold change of tumour size for fixed average treatment intensity and varied length of active phases/drug holidays.

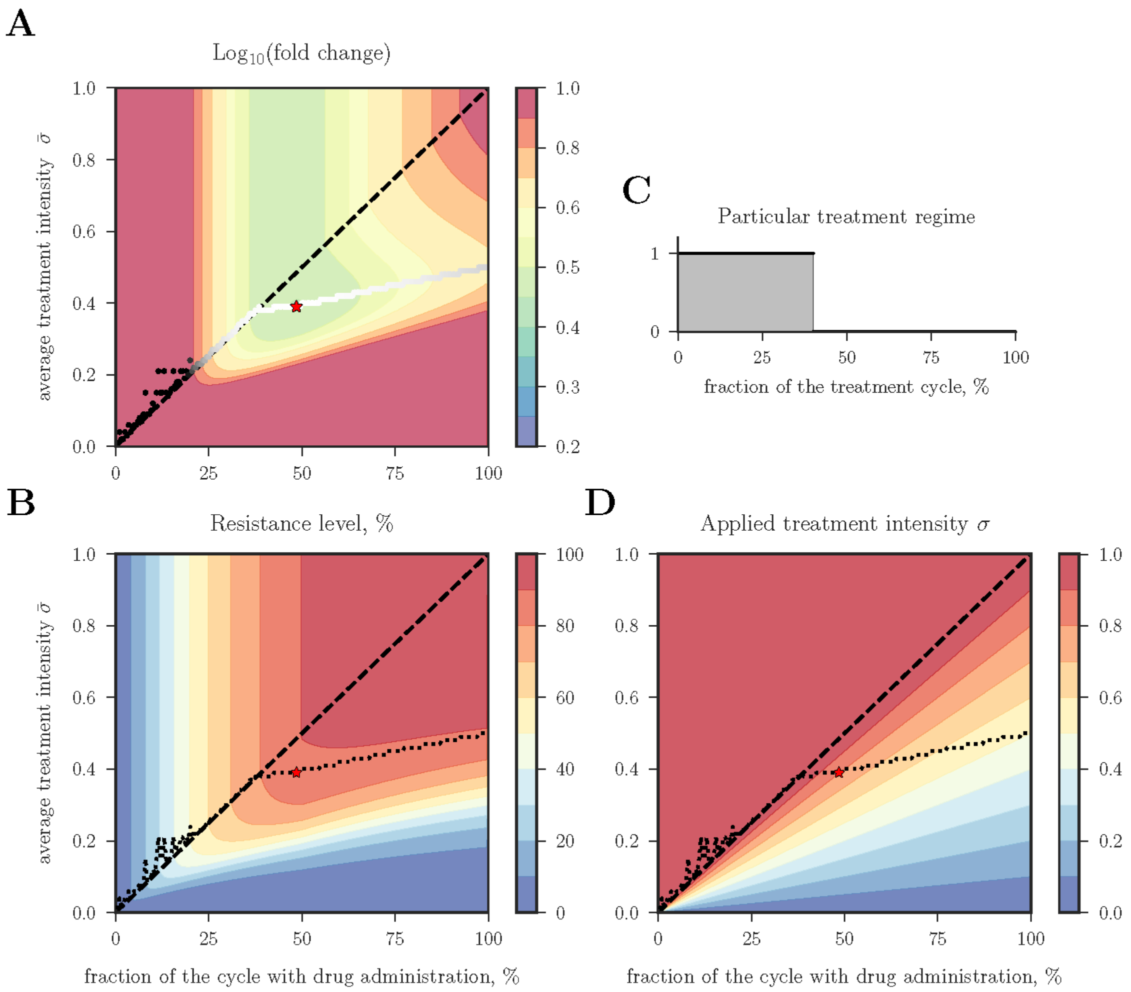


**Fig S5: Outcomes of the periodic treatment with 32-day periods of varying duty cycles (ratio of the active phase duration to the duration of the period): fold change in tumour size (A), final resistance level (B).** C and D illustrate the asymmetry in the treatment schedule and the applied treatment intensity during the active phase. The treatment intensity during the active phase is set to the MTD when it is larger than one (the region above the dashed diagonal in D). A line of dots shown in A, B and D indicates local minima, while the red star indicates the global minimum.

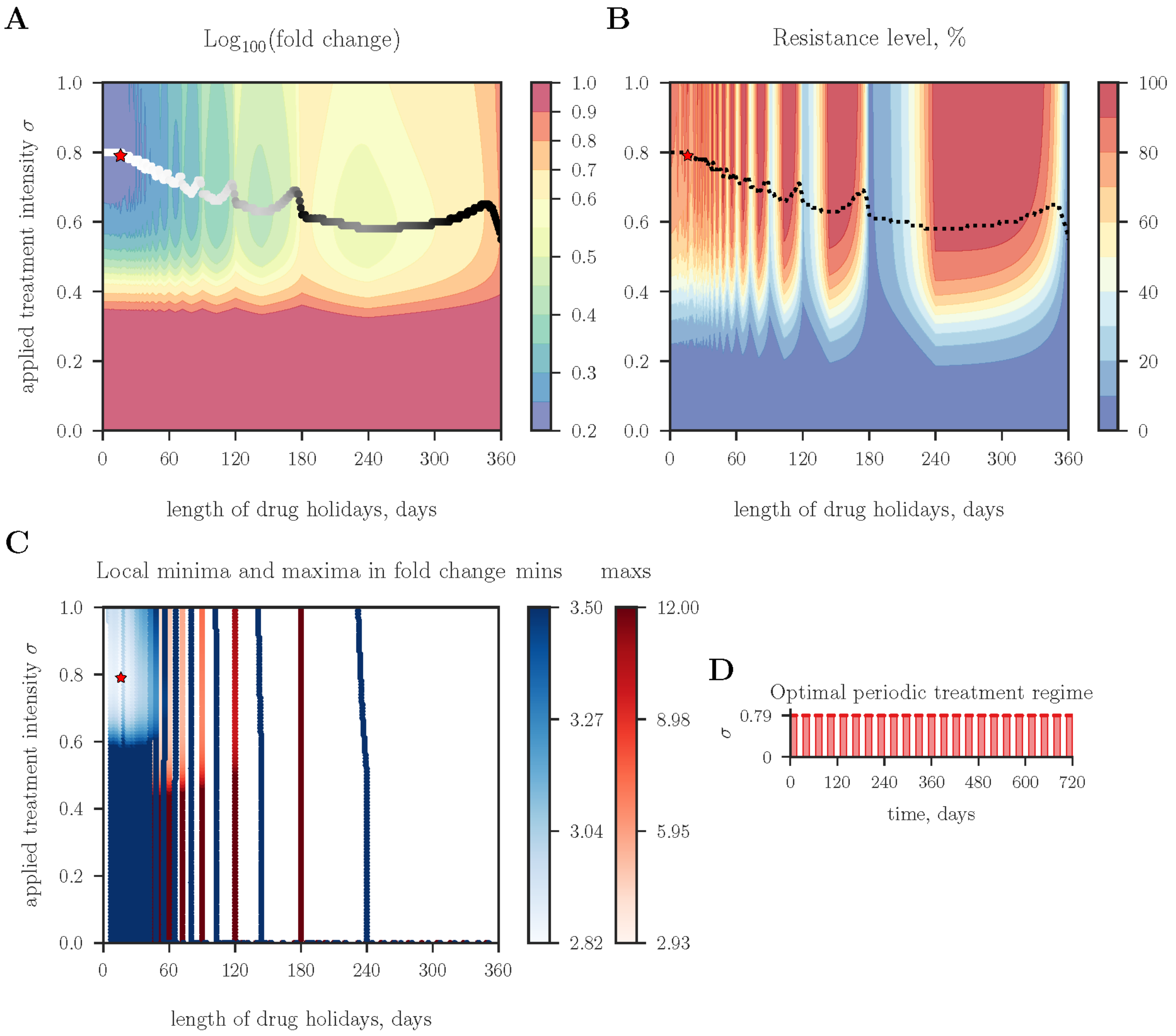


**Fig S6: Analogous to Fig S4 but with exception that all treatments are compared in terms of applied treatment intensity $\sigma$ rather than the average quantity $\bar{\sigma}$ (equivalent to cumulative treatment intensity), see *y*-axis.**

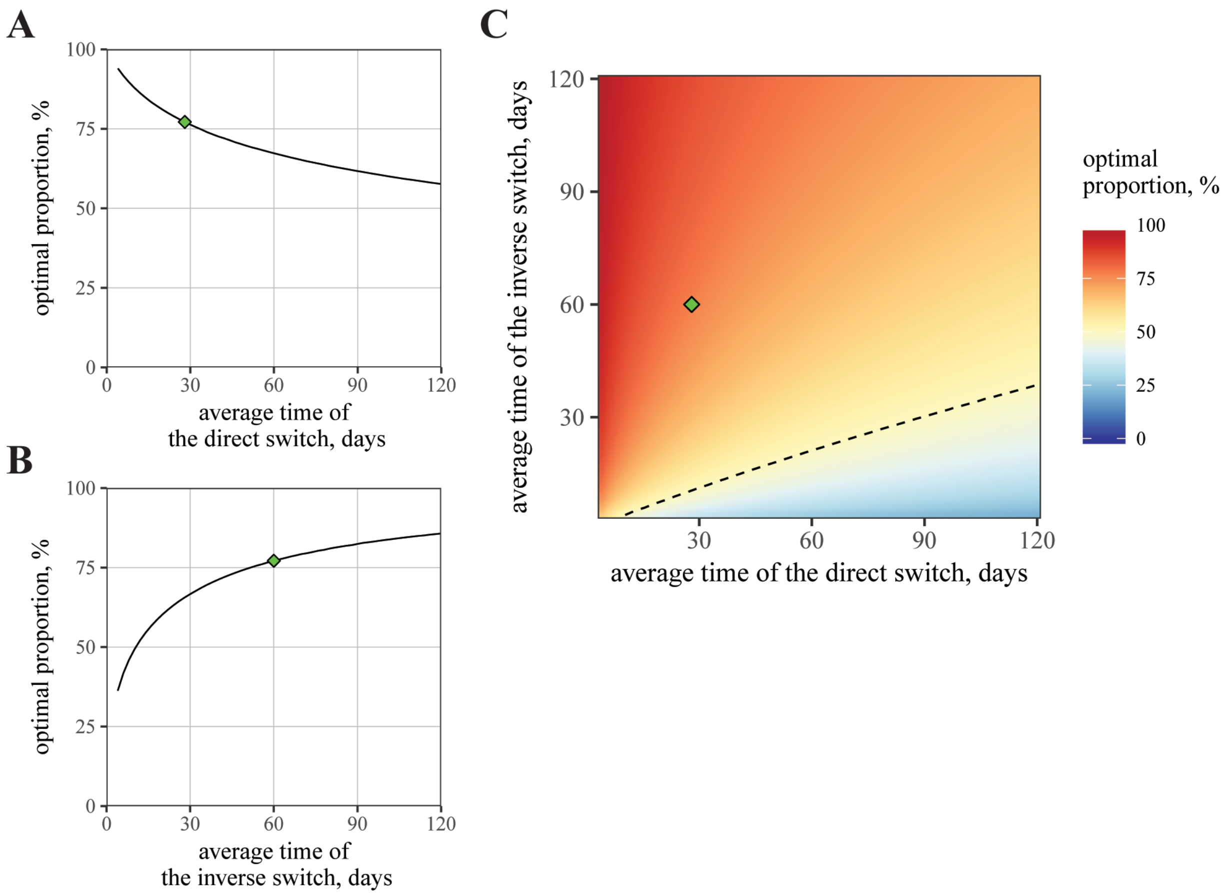


**Fig S7: Optimal balance between sensitive and resistant cells as a result of optimal problem.** (AB) The change in optimal proportion when one of switching rate is fixed at its baseline value (inverse switch at 60 days in A, direct switch at 28 days in B). (C) Variation in optimal proportion for varied characteristic times. Dashed line indicates the optimal proportion at 50%.

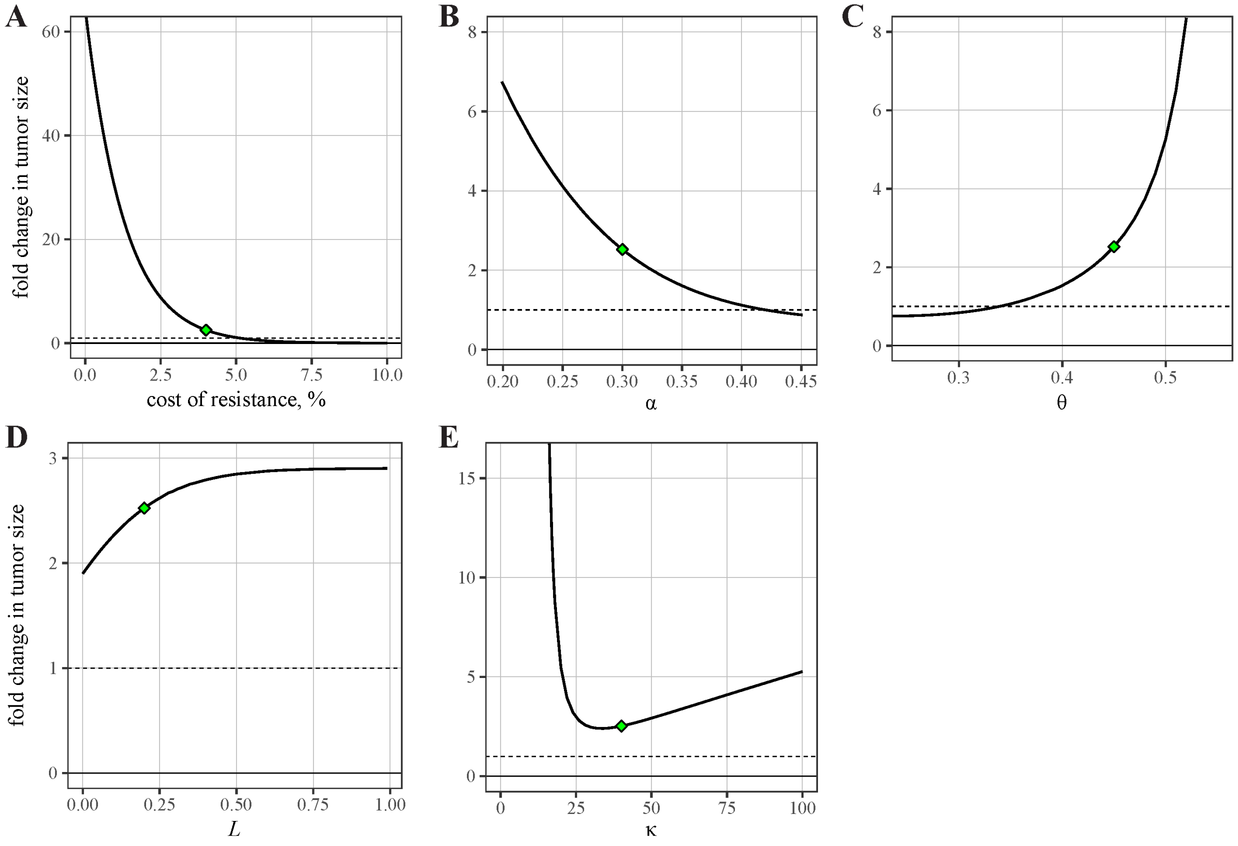


**Fig S8: Effects of single parameter values on the optimal outcome of the two-years treatment.** Other parameters remain fixed according to Table 1. The baseline parameters set and the corresponding outcome are shown as green points. The fold-change of level one is shown by dashed horizontal lines.

#### Appendix

A. Mathematical framework

We start with a model of antagonistic pathways for individual cells. We consider a stochastic birth-death process and apply the mean-field approximation to describe the evolution of tumour size and resistance level at any particular time of the treatment. We derive analytical equations for dynamics depending on the applied treatment intensity.

*A.1. Modelling pathway activity.*

We model a system of two interconnected pathways and an up-stream gene shown in Fig 1A. For simplicity, the main pathway is denoted by the index "1" and the alternative pathway by the index "2". We characterize each pathway with normalized activity level $y_i$ ($0 \le y_i \le 1$, $i = 1, 2$), and construct the protein expression model with two consistent reactions [1,2]. First, the production function $f_i(y_i)$ identifies the rate of protein production. Second, the degradation of proteins occurs at rate 1. Thus, the dynamics hold the form:

$$\rho^{-1}\frac{\mathrm{d}y_i(t)}{\mathrm{d}t} = f_i(y_i(t)) - y_{\mathrm{i}}(t) + \eta_i,$$

where $\eta_{1,2}$ are random noise terms, the constant $\rho$ describes a relative adjustment rate between updates in pathway activity and birth-death process of tumour cells. Following the accumulated experimental evidence of the cell fate to be guided by Boolean logic [3,4], we assign the production function $f_i(y_i)$ to a step-wise function: it equals to one for any $y_i$ above some threshold value $\theta_i$, and to $\alpha_i$ for any $y_i$ below it. This gives: $f_i(y_i) = \alpha_i + (1-\alpha_i)H(y_i - \theta_i)$, where $H(\cdot)$ is the Heaviside function, and $0 \le \alpha_i < \theta_i \le 1$ ($i = 1, 2$).

The activation dynamics can be equivalently described by the Brownian motion in a double-well potential: $U_i(y_i) = -\int f_i(y_i)\mathrm{d}y_i + y_i^2/2$. Two stable equilibria: up and down regulation states, correspond to the two minima of the potential (Fig 1D). The state $y_i$ fluctuating near the attractor $y^{\pm}$ may eventually jump from one potential well to another. The transition rate follows the Van't Hoff-Arrhenius law [5] and defines the escape rate from one of the equilibria $y_i^{\pm}$ in the form: $\lambda_i \exp(-\kappa_i E_i^{\pm})$, where $E_i^{\pm}$ are the heights of potential barriers. The constant $\lambda_i$ depends on the curvature of the potential in the proximity of the equilibrium and also at the boundary of the region of attraction $\theta_i \pm \varepsilon$ ($\varepsilon \to 0$).

We set the treatment intensity to the variable $\sigma$ ($0 \le \sigma \le 1$). The value $\sigma = 0$ indicates a situation of no treatment, while the value $\sigma = 1$ is used when the drug shuts down the main pathway completely. Intermediate values of $\sigma$ indicate relatively moderate levels of the treatment when the pathway activity is partially suppressed. We also notice that the treatment intensity $\sigma$ can be given by some non-linear (sigmoidal) function of the actual drug dosage. Additionally, the treatment can have a milder effect on the alternative pathway either due to imperfect targeting of the main pathway or chemotoxic side effect.

We model the drug action by modifying the production function:

$$f_i(y_i) = A_i(\sigma)\left[\alpha_i + (1-\alpha_i)H(y_i - \theta_i(\sigma))\right], \tag{S1}$$

where the function $A_i(\sigma)$ describes the reduction in expression level of the pathway, while the function $\theta_i(\sigma)$ translocates the threshold value $\theta_i$ (Fig S1A). We assume that $A_i(\sigma)$ decreases and $\theta_i(\sigma)$ shifts to the right with increasing $\sigma$. This shift imposes that more proteins are needed to maintain the pathway to be up-regulated.

We modify the potential accordingly:

$$U_i(y_i) = -f_i(y_i)(y_i - \theta_i(\sigma)) + \frac{y_i^2 - (\theta_i(\sigma))^2}{2}, \tag{S2}$$

and also impose that the potential $U_1(y_1)$ is a double well for any treatment intensity $\sigma < 1$, while it is unimodal only for $\sigma = 1$ (Fig S1B). The value $\sigma = 1$ relates to the maximally tolerated dosage (MTD). It has the equivalent effect on the dynamics as all other $\sigma > 1$, because the cell fitness is not affected directly by the drug action. Finally, we write the modified expression for the heights of potential barriers: $E_i^+ = U_i(\theta_i(\sigma)) - U_i(A_i(\sigma))$ and $E_i^- = U_i(\theta_i(\sigma)) - U_i(A_i(\sigma)\alpha_i)$.

Since the two pathways are mutually inhibitory, both pathways cannot be simultaneously activated. Hence, we present the whole dynamics of the system as a discrete dynamics of stochastic phase transitions. This means, for example, that a cell with active pathway 1 can transition to active pathway 2 only through a transient state, where both pathways are inactive (Fig 1B).

We call a cellular state with both pathways inactivated by “0”, with only main pathway activated by “1” (the proliferative state), with only alternative pathway activated by “2” (the quiescent state). Then we define the probability $P_k(t)$ that a cell randomly sampled from the whole population of tumour cells is at state $k = 0,1,2$. The dynamics are written as follows:

$$\frac{dP_i(t)}{dt} = -P_i(t)\lambda_i e^{-\kappa_i E_i^+} + P_0(t)\lambda_i e^{-\kappa_i E_i^-},$$

where $i = 1,2$ and $P_0(t) = 1 - P_1(t) - P_2(t)$.

*A.2. Tumour growth as a stochastic birth-death process.*

We further describe the dynamics of tumour growth at population level by taking into account the process of aforementioned individual pathway activities.

We set a characteristic time period between two consecutive cell divisions to a variable $\Delta T$ (order of 3–7 days for cancer cells [6,7]). First, we elaborate on a discrete-time version for the birth-death process. The relative number of cells of type $k$ ($k = \{0,1,2\}$, Fig 1B) obeys the dynamics: $n_k\big((m+1)\Delta T\big) = P_k\big((m+1)\Delta T\big) n(m\Delta T)(1+\omega_k)$, for the current time step $m = 1,2,\ldots$; $P_k((m+1)\Delta T)$ is obtained by solving Equation (S1) on the time interval $[m\Delta T, (m+1)\Delta T]$ and with initial conditions given at time $t = m\Delta T$: $P_k(m\Delta T) = n_k(m\Delta T)/n(m\Delta T)$ ($n(t) = \sum_k n_k(t)$ is a tumour load). The variable $\omega_k$ denotes the difference between birth and death rates, i.e. the fitness of a cell with state $k$. In case when the alternative pathway is activated, we incur the fitness cost $c$, such that: $\omega_2 = \omega_1 - c$. Thus, the following relation holds: $\omega_0 < 0 < \omega_2 < \omega_1$. The first inequality in the chain reflects the fact that the cell becomes nonviable when both pathways are shut down.

To write the continuous-time version of the dynamics, we study a limiting case when the length of the time step becomes arbitrarily small. In that case we derive the following equations:

$$\frac{\mathrm{d}n_k(t)}{\mathrm{d}t} = \omega_k n_k(t) + \mu \frac{\mathrm{d}P_k(t)}{\mathrm{d}t} n(t),$$

where $\mu$ is the rate of updates for pathway activity.

To observe the characteristic *U*-shape curve in tumour dynamics under treatment application, when the initial tumour remission changes to final stage of tumour relapse, see e.g. [8], here we require the update rates in pathway activity to be moderate ($\mu \sim 1$).

We re-write (S1) in the form:

$$\frac{1}{\lambda_i e^{-\kappa_i E_i^-}} \frac{\mathrm{d}P_i(t)}{\mathrm{d}t} = 1 - P_i(t)(1 + e^{-\kappa_i \Delta E_i}) - P_{i'}(t),$$

where $\Delta E_i = E_i^+ - E_i^-$, $i'$ is complementary to $i$ ($i' = 3 - i$ for $i = \{1,2\}$). Substituting it to the previous equation for the dynamics of $n_k(t)$, we derive the following equations after some algebra:

$$\frac{\mathrm{d}x_i(t)}{\mathrm{d}t} = (\omega_i - \bar{\omega})x_i(t) + \mu_i e^{-\kappa_i E_i^-}(x_0(t) - x_i(t)e^{-\kappa_i \Delta E_i}),$$

$$\frac{\mathrm{d}x_0(t)}{\mathrm{d}t} = (\omega_0 - \bar{\omega})x_0(t) - \sum_{i=1}^{2} \mu_i e^{-\kappa_i E_i^-}(x_0(t) - x_i(t)e^{-\kappa_i \Delta E_i}),$$

where $x_k(t) = n_k(t) / n(t)$ are relative frequencies of cells of type $k = \{0,1,2\}$. Whereas two parameters $\mu_1$ and $\mu_2$ are expressed through the rates $\mu$, $\lambda_1$ and $\lambda_2$, and introduced for convenience. The parameter $\bar{\omega}$ is chosen to guarantee the identity: $\sum_k x_k(t) = 1$. Namely, it equals to the average fitness value over the whole population: $\bar{\omega} = \sum_k \omega_k x_k(t)$ [9].

We do the following transformation of state variables [10]:

$$\xi_k(0) = x_k(0), \quad \xi_k(t) = x_k(t)\exp\left(\int_0^t \bar{\omega}\mathrm{d}t\right), \tag{S3}$$

to convert the previous equations to the linear form :

$$\begin{aligned} \frac{\mathrm{d}\xi_i(t)}{\mathrm{d}t} &= \omega_i \xi_i(t) + \mu_i e^{-\kappa_i E_i^-}(\xi_0(t) - \xi_i(t) e^{-\kappa_i \Delta E_i}), \\ \frac{\mathrm{d}\xi_0(t)}{\mathrm{d}t} &= \omega_0 \xi_0(t) - \sum_{i=1}^{2} \mu_i e^{-\kappa_i E_i}(\xi_0(t) - \xi_i(t) e^{-\kappa_i \Delta E_i}). \end{aligned} \tag{S4}$$

Finally, we describe the dynamics of a tumour load $n(t)$ by using the transformation (S3): $\mathrm{d}n(t) / \mathrm{d}t = \bar{\omega} n(t)$, and then:

$$n(t) = n(0)\exp\left(\int_0^t \bar{\omega}\mathrm{d}t\right) = n(0)\sum_{k=0}^{2} \xi_k(t).$$

*A.3. Implementation of the fast- and slow-time dynamics in the pathway activity.*

To account for biological evidence, we introduce two different time-scales in the model. We assume that any arrangement in protein expression numbers within a given (proliferative or quiescent) cellular state – i.e. the transition rates between states "0" and "1" or between "0" and "2" – occurs arbitrarily fast. However, the transition $1 \to 0 \to 2$ and vice versa implies the change in the cellular phenotype, so it requires not only the alteration of pathway activities, but also some physiological changes of cellular phenotypes. So, we implement that this change occurs at a much slower rate.

Thus, we encode a cellular state by two values: $k_a$, where $k$ is the current state and a is the previous state ($k \neq a$; $k, a = \{0,1,2\}$; Fig 1C). Analogously, the state $1_0$ denotes the proliferative cells with up-regulated pathway 1, the state $2_0$ denotes the quiescent resistant cells with up-regulated pathway 2. Whereas the state $0_1$ implies the transients state of proliferative cells with low activity on both pathways, the state $0_2$ implies the transient state of quiescent cells with low activities on both pathways. The transition dynamics $k_0 \leftrightarrow 0_k$ occurs at fast-time scale, so that the equilibrium in corresponding subsystems (red and grey compartments in Fig 1C) is achieved instantaneously. In contrast, the transition $0_{i'} \to i_0$ occurs at a slow scale with rates $\mu_i \sim 1$ ($i' = 3-i$, $i = \{1,2\}$).

Thus, we derive a modified version of the dynamics (S4). As before, we introduce new variables $\xi_{ka}$. The fast-time dynamics give:

$$\xi_{0i}(t) = \xi_{i0}(t) e^{-\kappa_i \Delta E_i}, \quad i = \{1,2\}, \tag{S5}$$

so that we have:

$$\begin{aligned} \frac{\mathrm{d}\xi_{i0}(t)}{\mathrm{d}t} &= \omega_i \xi_{i0}(t) + \mu_{i'} e^{-\kappa_i E_i^-} \xi_{0i'}(t), \\ \frac{\mathrm{d}\xi_{0i}(t)}{\mathrm{d}t} &= \omega_0 \xi_{0i}(t) + \mu_i e^{-\kappa_{i'} E_{i'}^-} \xi_{0i}(t), \end{aligned} \tag{S6}$$

where $i'$ is the complement to $i$ ($i' = 3-i$). Then we claim that both state variables $\xi_{i0}$ and $\xi_{0i}$ describe the same phenotype of the cell $i$, so that we introduce a new variable: $\bar{\xi}_i(t) = \xi_{i0}(t) + \xi_{0i}(t)$, to obtain from (S5)–(S6):

$$\frac{\mathrm{d}\bar{\xi}_i(t)}{\mathrm{d}t} = \frac{\omega_i e^{\kappa_i \Delta E_i} + \omega_0}{e^{\kappa_i \Delta E_i} + 1} \bar{\xi}_i(t) - \frac{\mu_i e^{-\kappa_{i'} E_{i'}^-}}{e^{\kappa_i \Delta E_i} + 1} \bar{\xi}_i(t) + \frac{\mu_{i'} e^{-\kappa_i E_i^-}}{e^{\kappa_{i'} \Delta E_{i'}} + 1} \bar{\xi}_{i'}(t). \tag{S7}$$

If we apply additionally: $\omega_i = b(1 - c(i-1)) - d$ and $\omega_0 = b(1-\chi) - d$, where $b$ and $d$ are birth and death rates respectively, $c$ is the (additive) cost of resistance, $\chi$ is the penalty to the birth rate due to knocking down the cellular machinery, we get the following expression for the fitness of a cell of type $i$:

$$\bar{\omega}_i = \frac{\omega_i e^{\kappa_i \Delta E_i} + \omega_0}{e^{\kappa_i \Delta E_i} + 1} = b \frac{1 + e^{\kappa_i \Delta E_i} - \chi - c e^{\kappa_i \Delta E_i}(i-1)}{e^{\kappa_i \Delta E_i} + 1} - d, \tag{S8}$$

that represents the first term in (S7).

From now on, we consider only the case when the treatment targets the main pathway ideally. This imposes no effect of the treatment on the cells with active alternative pathway. Using (S7)–(S8), we write:

$$\begin{aligned} \frac{\mathrm{d}\bar{\xi}_1(t)}{\mathrm{d}t} &= \left( \frac{b e^{\kappa_1 \Delta E_1} + b(1-\chi) - \mu_1}{e^{\kappa_1 \Delta E_1} + 1} - d \right) \bar{\xi}_1(t) + \mu_2 e^{-\kappa_1 E_1^-} \bar{\xi}_2(t), \\ \frac{\mathrm{d}\bar{\xi}_2(t)}{\mathrm{d}t} &= (b(1-c) - \mu_2 e^{-\kappa_1 E_1^-} - d) \bar{\xi}_2(t) + \frac{\mu_1 \bar{\xi}_1(t)}{e^{\kappa_1 \Delta E_1} + 1}, \end{aligned} \tag{S9}$$

where we re-define certain parameters $c$, $\mu_1$, $\mu_2$ for our convenience. Notice that $\Delta E_1$ and $E_1^-$ are functions of the treatment intensity $\sigma$.

Dynamics (S9) allow to derive an analytical formula for evolution of relative frequency for resistant cells that we denote by $x(t)$: $x_1(t) \equiv 1 - x(t)$ and $x_2(t) \equiv x(t)$. By definition (S3), we have: $x(t) = \bar{\xi}_2(t) / (\bar{\xi}_1(t) + \bar{\xi}_2(t))$ that leads to dynamical equations (1) and (2) in the main text, where we also omit the subindices $\kappa \equiv \kappa_1$, $E^- \equiv E_1^-$ and $\Delta E \equiv \Delta E_1$, additionally with $\mu \equiv \mu_1$ and $\bar{\mu} \equiv \mu_2$ for short hand:

$$\frac{\mathrm{d}n(t)}{\mathrm{d}t} = \left( b\left(1 - \frac{\chi(1 - x(t))}{e^{\kappa\Delta E} + 1} - cx(t)\right) - d \right) n(t), \tag{10}$$

$$\frac{\mathrm{d}x(t)}{\mathrm{d}t} = b\left(\frac{\chi}{e^{\kappa\Delta E} + 1} - c\right) x(t)(1 - x(t)) + \frac{\mu(1 - x(t))}{e^{\kappa\Delta E} + 1} - \bar{\mu}x(t). \tag{11}$$

The main terms of **Error! Reference source not found.**–(11): $(e^{\kappa\Delta E} + 1)$ and $e^{-\kappa E^-}$, are monotonic decreasing functions of the treatment intensity σ (Fig S2). Thus, the resistant cells overgrow sensitive cells for any value $\mu$ and $\bar{\mu}$ only if $\chi / \left(e^{\kappa\Delta E} + 1\right) > c$, which is expected for large values of drug dosages $\sigma$.

*A.4. Model simulation and determination of model parameter values*

We simulate tumour growth with or without drug administration by solving the aforementioned dynamical equations. To make the simulation outcomes comparable to real data, we estimate and extract model parameters from experimental literature. Below we list the source and criteria for determining model parameters.

Bozic *et al.* [7] reported the average daily net growth rate of a melanoma cell-line with BRAF mutated gene at 0.01. They set the death rate to 0.13 per day, while the birth rate was set to 0.14 per day to ensure the cells to divide every 7 days on average. The treatment with BRAF inhibitor *vemurafenib* led to the observed tumour shrinkage at average rate of 0.03 per day.

If the action of the drug has a weak cytotoxic effect, there is no change in the death rate, while the therapeutic agent disrupts the internal cellular machinery making a cell unable to divide. Additionally, assuming that Bozic *et al.* used the drug intensity close to maximally tolerated dosage (MTD), we arrive at the following equations that are particular cases of the dynamics (1):

$$b\left(1 - \frac{\chi}{e^{\kappa\Delta E(\sigma=0)} + 1}\right) - d = 0.01\,\mathrm{day}^{-1}, \qquad b\left(1 - \frac{\chi}{e^{\kappa\Delta E(\sigma=1)} + 1}\right) - d = -0.03\,\mathrm{day}^{-1},$$

where $d = 0.13$ day$^{-1}$. This in turn leads to the following conditions:

$$b = \frac{0.1(e^{\kappa\Delta E(\sigma=1)}+1) - 0.14(e^{\kappa\Delta E(\sigma=0)}+1)}{e^{\kappa\Delta E(\sigma=1)} - e^{\kappa\Delta E(\sigma=0)}}, \qquad \chi = 1 - \frac{0.14(e^{\kappa\Delta E(\sigma=0)}+1) - be^{\kappa\Delta E(\sigma=0)}}{b}.$$

From the fixed parameters characterizing the dynamics of the main pathway activity: expression level at the down state $\alpha = 0.3$, threshold value for the production function $\theta = 0.45$, robustness parameter $\kappa = 40.0$, and a parameter characterizing the effect of the drug to shift the threshold $\theta$ in the production function $L = 0.2$ (see Table 1 for parameter description), we obtain the following fitted values $b = 0.14$ day$^{-1}$ and $\chi = 0.30$. The characteristic time of switching from the main to the alternative pathway is set to be 28 days, and of the reversed switch to be 60 days. The former number is chosen by assuming that rewiring of cellular machinery requires at least several cell divisions and not only instant change in pathway activity (e.g. experimental evidence for a tumour treated with kinase inhibitors, see [11,12]). The latter value of 60 days is taken from assumption that the reverse switch occurs either much slower comparing to the direct switch or remains irreversible in some cases [13]. The switching rate is the inverse quantity of the characteristic time. Resistant cells have a positive fitness advantage only if the cost of resistance does not exceed the threshold value of $1 - d/b = 13\%$. Above that value, a population of cells with the activated alternative pathway is not sustainable for any treatment. Thus, we set the cost of resistance in our simulation to approximately one third of the threshold value $c = 4\%$. Since the proposed model parameter values are obtained from either educated guess or similar theoretical studies [1,2,14], we also conduct sensitivity analysis on simulation outcomes.

B. Design of the optimal treatment.

*B.1. Formulation of the optimal control problem.*

Here we use a well-known technique from the Optimal Control Theory to design the optimal treatment regimen that minimizes the tumour size after a fixed time period. Mathematically, this leads to a solution of the first order partial differential equation that is also known as Hamilton-Jacobi-Bellman (HJB) equation. The unknown function represents the expected outcome of the treatment starting from any given state of the system $(t, n(t), x(t))$. That quantity is usually referred as the value function. Specifically for our problem, we define the value function as the logarithm of the tumour fold change at the terminal time $T$ that has the following form:

$$\ln\left(\frac{n(T)}{n(0)}\right) = \int_0^T \left[ b\left(1 - \frac{\chi(1-x(t))}{e^{\kappa\Delta E}+1} - cx(t)\right) - d \right] \mathrm{d}t \to \min_{\sigma\in[0,1]}.$$

We can re-write it in shorter (and more convenient) form that neglects the constants in the integral and changes the sign in front of the integral:

$$\int_0^T \left( \frac{\chi(1-x(t))}{e^{\kappa\Delta E}+1} + cx(t) \right) \mathrm{d}t \to \max_{\sigma\in[0,1]}.$$

Finally, we define the HJB equation as follows:

$$\mathcal{H} \doteq -\phi_0 + \max_{\sigma\in[0,1]}\left[\phi\left(b\left(\frac{\chi}{e^{\kappa\Delta E}+1}-c\right)x(1-x)+\frac{\mu(1-x)}{e^{\kappa\Delta E}+1}-\bar{\mu}e^{\kappa E^{-}}x\right)+\frac{\chi(1-x)}{e^{\kappa\Delta E}+1}+cx\right]=0, \quad \text{(S12)}$$

subject to the dynamics (1)–(2) in the main text. Here we introduced two costate variables denoted by variables $\phi_0$ and $\phi$. They provide the components of the gradient function of the value function. By definition, $\phi(T)=0$. Since this condition for $\phi$ is given at the terminal time $T$, it will be convenient to follow the dynamics in backward time $\tau = T - t$.

To construct the field of optimal trajectories, we emit the characteristics associated with solution of the HJB equation (S12). Each characteristic represents a solution of the system of characteristics, a system of first order ordinary differential equations, see e.g. [15] for details. Precisely speaking, one of the characteristic equations is already given by the dynamics (2) in the main text:

$$\frac{\mathrm{d}x(\tau)}{\mathrm{d}\tau} = -\frac{\partial\mathcal{H}}{\partial\phi} = -b\left(\frac{\chi}{e^{\kappa\Delta E}+1}-c\right)x(\tau)(1-x(\tau)) - \frac{\mu(1-x(\tau))}{e^{\kappa\Delta E}+1} + \bar{\mu}x(\tau).$$

Another equation for $\phi(t)$ is obtained by differentiating the Hamiltonian (S12) with respect to $x$:

$$\frac{\mathrm{d}\phi(\tau)}{\mathrm{d}\tau} = \frac{\partial\mathcal{H}}{\partial x} = \phi(\tau)\left(b\left(\frac{\chi}{e^{\kappa\Delta E}+1}-c\right)(1-2x(\tau)) - \frac{\mu}{e^{\kappa\Delta E}+1} - \bar{\mu}e^{-\kappa E^{-}}\right) - \frac{\chi}{e^{\kappa\Delta E}+1} + c.$$

We finally complement these two equations with the expression to define the optimal treatment at each particular time. It is derived from maximizing (S12):

$$\sigma^{*} = \arg\max_{\sigma\in[0,1]}\mathcal{H} = \arg\max_{\sigma\in[0,1]}\left(\frac{\phi(b\chi x+\mu)+\chi}{e^{\kappa\Delta E}+1}(1-x) - \bar{\mu}\phi e^{-\kappa E^{-}}x\right), \quad \text{(S13)}$$

where the expression in brackets is a collection of all terms in the Hamiltonian $\mathcal{H}$ that depend on $\sigma$. The value $\sigma^{*}$ equals to zero or one, if the maximum of the Hamiltonian is achieved at one of the boundaries of the segment [0,1]. Otherwise, $\sigma^{*}$ is assigned to the intermediate value of [0,1] at which $\partial\mathcal{H}/\partial\sigma|_{\sigma=\sigma^{*}} = 0$.

The treatment intensity can be obtained at the terminal time:

$$\sigma^*(T) = \arg\max_{\sigma\in[0,1]} (\chi(1-x(T))/(e^{\kappa\Delta E}+1)) = 1.$$

In the following time moments, it requires the knowledge of the state and costate variables $(x(t), \phi(t))$.

*B.2. Primary field of optimal trajectories.*

To proceed further, we need to study the behaviour of the function in brackets in (S13). It is composed of two sub-functions:

$$\sigma^* = \arg\max_{\sigma\in[0,1]} \rho(\sigma), \quad \text{where} \quad \rho(\sigma) = \rho_1(\sigma) + \rho_2(\sigma),$$

and the first sub-function: $\rho_1(\sigma) = (\phi(b\chi x + \mu) + \chi)(1-x)/(e^{\kappa\Delta E}+1)$, monotonically increasing for any $\tau \geq 0$, second sub-function: $\rho_2(\sigma) = -\bar{\mu}\phi e^{-\kappa E^-} x$, monotonically decreasing for any $\tau > 0$ and equal exactly zero at $\tau = 0$ ($\tau = T - t$). This means that at the terminal time the global maximum is reached at $\sigma = 1$ (Appendix Fig 1A). Whereas for further moments $\tau > 0$, the function $\rho_2(\sigma)$ start to elevate for $\sigma \in [0,1]$, so that the global maximum may start drifting from $\sigma = 1$. At first, the elevation of $\rho_2$ leads to appearance of the intermediate maximum where $0 < \sigma^* < 1$ (Appendix Fig 1B). For some trajectories with $x(T) > \bar{x}$, the function $h_2$ goes even higher, so that the global maximum switches to $\sigma^* = 0$ (Appendix Fig 1C). For other trajectories with $x(T) < \bar{x}$, this does not occur and the value of $h_2$ at $\sigma = 0$ stays lower than the intermediate maximum of $h_1(\sigma^*) + h_2(\sigma^*)$ at some $0 < \sigma^* < 1$. Such observations result in construction of the primary field of trajectories shown in blue, and the switching curve $\mathcal{S}_1$, shown in dashed in Appendix Fig 2.

This completes the first step for constructing the optimal trajectories emitted directly from the terminal surface $t = T$. We identify such threshold value $\bar{x}$ that all trajectories emitted from the points below ($0 \leq x(T) < \bar{x}$) yield the active drug administration for the whole time period ($\sigma^* > 0$ at any $\tau \geq 0$); while all trajectories emitted from the points above ($\bar{x} < x(T) \leq 1$) contain an abrupt switch in the treatment (the patient is transferred from a non-zero drug dosage to a no-treatment stage at some given moment of time). The switching times are varied depending on $x(T)$ and form the switching curve in the state space that is indicated as $\mathcal{S}_1$ in Appendix Fig 2.

*B.3. Final steps of the solution.*

However, a part of the state space remains still uncovered by the characteristics. Indeed, the monotonicity of the trajectory $x(\tau)$ emitted from a terminal point slightly above $\bar{x}$ ($x(T) = \bar{x} + \varepsilon$, where $\varepsilon \to +0$) changes from decreasing to increasing at the moment of switch. In contrast, it does not occur for the trajectory emitted from a terminal point slightly below $\bar{x}$ ($x(T) = \bar{x} - \varepsilon$). This region is covered by orange trajectories in Appendix Fig 2, and our next aim will be to identify the method on how to cover the empty space with trajectories, and, namely, how to construct the trajectories in orange and a special curve $\mathcal{S}_2$.

To cover the whole state space with optimal trajectories, we emit a special (singular) trajectory from the ending point of the switching curve. This represents a usual resolution present in problems in theory of optimal control with such curve known under the name of a singular arc in theory of linear control [15], or a universal singular characteristic in non-linear control [16,17].

The singular curve is characterized by non-smoothness of the value function and can be constructed using the method of singular characteristics, see [15,18]. The resulting control is of the chattering type, consisting of on'n'off phases with vanishingly short time lengths. Intuitively, we seek for such curve that obeys two possible candidates for the maximizing function of $\sigma^*$ such as shown in Appendix Fig 1D. To define that curve, we introduce a so-called singular Hamiltonian that can be written in its general form as follows:

$$\nu\mathcal{H}^{\text{sing}} \doteq \{\mathcal{H}_1,\mathcal{H}_0\}\mathcal{H}_{-1} + \{\mathcal{H}_0,\mathcal{H}_{-1}\}\mathcal{H}_1 + \{\mathcal{H}_{-1},\mathcal{H}_1\}\mathcal{H}_0,$$

with three necessary conditions hold on the curve: $\mathcal{H}_1 = 0$, $\mathcal{H}_0 = 0$, and $\mathcal{H}_{-1} = 0$, while $\nu$ representing a scaling parameter. The curly brackets denote the Poisson brackets that writes for two Hamiltonians $\mathcal{F}$ and $\mathcal{G}$ as:

$$\{\mathcal{F},\mathcal{G}\} \doteq \frac{\partial\mathcal{F}}{\partial x}\times\frac{\partial\mathcal{G}}{\partial\phi} - \frac{\partial\mathcal{F}}{\partial\phi}\times\frac{\partial\mathcal{G}}{\partial x},$$

and can be easily calculated once $\mathcal{F}$ and $\mathcal{G}$ are given. Then the singular curve satisfies the solution of the characteristic system that is similar to the ordinary one, but with the singular Hamiltonian $\mathcal{H}^{\text{sing}}$ instead of ordinary Hamiltonian $\mathcal{H}$:

$$\frac{\mathrm{d}x(\tau)}{\mathrm{d}\tau} = -\nu\frac{\partial\mathcal{H}^{\text{sing}}}{\partial\phi}, \quad \frac{\mathrm{d}\phi(\tau)}{\mathrm{d}\tau} = \nu\frac{\partial\mathcal{H}^{\text{sing}}}{\partial x}.$$

In our case we have: $\mathcal{H}_1 \equiv \mathcal{H}$, $\mathcal{H}_0 \equiv \mathcal{H}|_{\sigma=0}$, and $\mathcal{H}_{-1} \equiv \{\mathcal{H}_1,\mathcal{H}_0\}$. By following the representation adopted from [16]: $\nu = \gamma_0 + \gamma_1$ with $\gamma_0 = \{\{\mathcal{H},\mathcal{H}_0\},\mathcal{H}\}$ and $\gamma_1 = \{\{\mathcal{H}_0,\mathcal{H}\},\mathcal{H}_0\}$, we obtain the characteristic equations in the form:

$$\frac{\mathrm{d}x(\tau)}{\mathrm{d}\tau} = -\frac{1}{\gamma_0+\gamma_1}\left(\gamma_0\frac{\partial\mathcal{H}_0}{\partial\phi} + \gamma_1\frac{\partial\mathcal{H}}{\partial\phi}\right), \quad \frac{\mathrm{d}\phi(\tau)}{\mathrm{d}\tau} = \frac{1}{\gamma_0+\gamma_1}\left(\gamma_0\frac{\partial\mathcal{H}_0}{\partial x} + \gamma_1\frac{\partial\mathcal{H}}{\partial x}\right).$$

Complemented with the boundary condition given at the end of the switching curve $\mathcal{S}_1$ (Appendix Fig 1), we obtain the solution numerically and construct the singular curve.

As the next step, we emit two characteristic field of trajectories: one above the singular curve with $\sigma^* > 0$ and another below the singular curve with $\sigma^* = 0$. They cover the remaining part of the phase space with the trajectories that are shown in orange in Appendix Fig 1.

Finally, we verify that the obtained solution indeed obeys the consistency property when any initial condition determines a unique optimal trajectory. It also satisfies all necessary conditions to be a viscous solution of the Hamilton-Jacobi-Bellman equation. The latter is required by the mathematical theory of the first order partial differential equations [19].

Appendix figures

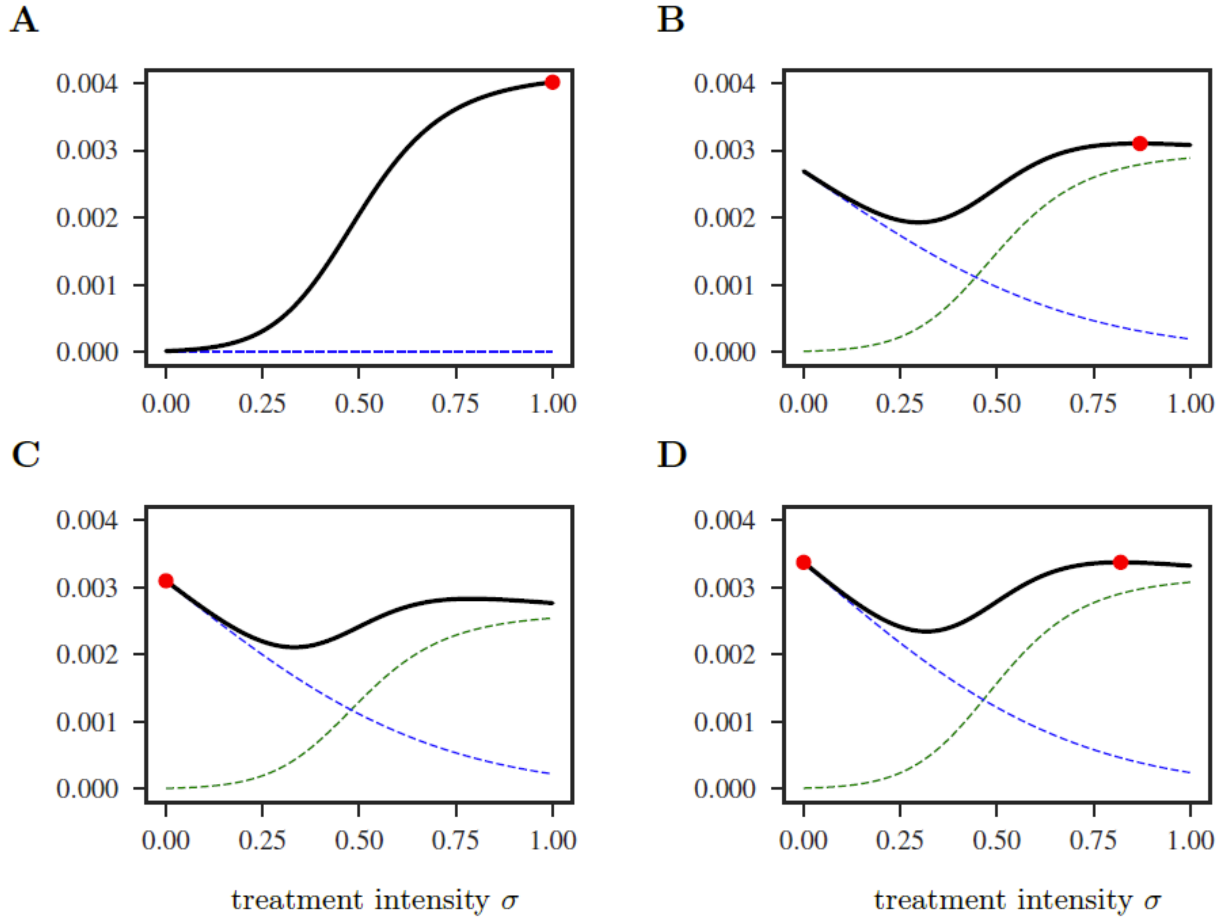


**Appendix Fig 1: Behaviour of the function $\rho(\sigma)$ depending on the point within the phase space $(\tau, x, \phi)$.** Each subplot shows different states: (A) on the terminal surface with $x(T) = 0.9$; (B) in the primary field of the trajectory prior the switching moment with $\tau = 12.24$, $x = 0.86$, $\phi = -0.29$, and (C) after the switching moment with $\tau = 28.8$, $x = 0.85$, $\phi = -0.34$; (D) on the singular characteristic (universal line) with $\tau = 50.05$, $x = 0.77$, $\phi = -0.41$. The sub-functions $\rho_1(\sigma)$ and $\rho_2(\sigma)$ are in dashed green and dashed blue respectively. Red dots denote the position of the global maxima, therefore indicating the value of optimal control $\sigma^*$. Parameter values are as Table 1.

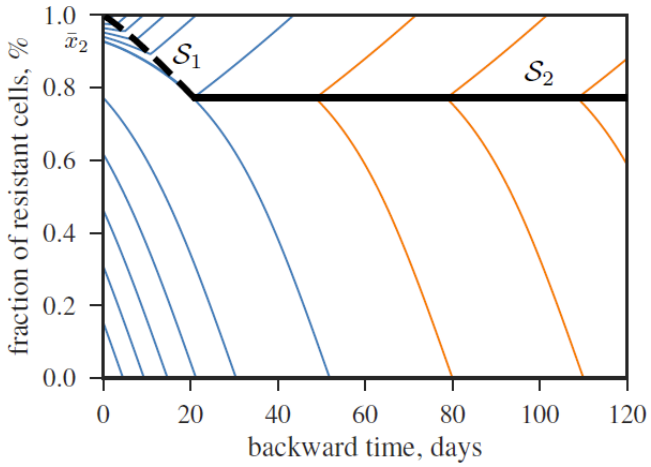


**Appendix Fig 2: Pattern of optimal trajectories is characterized by a switching curve $\mathcal{S}_1$ (black dashed) and a singular curve of universal type $\mathcal{S}_2$ (black solid).** Primary field of trajectories emitted from the terminal time consists of blue curves. While the rest of trajectories shown in orange are constructed using the curve $\mathcal{S}_2$. The threshold resistance level $\bar{x}_2$ separates the trajectories with and without a switch. Model parameters as in Table 1.

1. Assaf M, Roberts E, Luthey-Schulten Z, Goldenfeld N. 2013 Extrinsic Noise Driven Phenotype Switching in a Self-Regulating Gene. *Phys. Rev. Lett.* **111**. (doi:10.1103/PhysRevLett.111.058102)

2. Roberts E, Be'er S, Bohrer C, Sharma R, Assaf M. 2015 Dynamics of simple gene-network motifs subject to extrinsic fluctuations. *Phys. Rev. E* **92**. (doi:10.1103/PhysRevE.92.062717)

3. Bernardo-Faura M, Massen S, Falk CS, Brady NR, Eils R. 2014 Data-Derived Modeling Characterizes Plasticity of MAPK Signaling in Melanoma. *PLoS Comput. Biol.* **10**, e1003795. (doi:10.1371/journal.pcbi.1003795)

4. Olsson A, Venkatasubramanian M, Chaudhri VK, Aronow BJ, Salomonis N, Singh H, Grimes HL. 2016 Single-cell analysis of mixed-lineage states leading to a binary cell fate choice. *Nature* **537**, 698–702. (doi:10.1038/nature19348)

5. Hänggi P, Talkner P, Borkovec M. 1990 Reaction-rate theory: fifty years after Kramers. *Rev. Mod. Phys.* **62**, 251–341. (doi:10.1103/RevModPhys.62.251)

6. Bozic I *et al.* 2010 Accumulation of driver and passenger mutations during tumor progression. *Proc. Natl. Acad. Sci.* **107**, 18545–18550. (doi:10.1073/pnas.1010978107)

7. Bozic I *et al.* 2013 Evolutionary dynamics of cancer in response to targeted combination therapy. *eLife* **2**. (doi:10.7554/eLife.00747)

8. Chmielecki J *et al.* 2011 Optimization of Dosing for EGFR-Mutant Non-Small Cell Lung Cancer with Evolutionary Cancer Modeling. *Sci. Transl. Med.* **3**, 90ra59-90ra59. (doi:10.1126/scitranslmed.3002356)

9. Nowak MA. 2006 *Evolutionary dynamics: exploring the equations of life*. Cambridge, Mass: Belknap Press of Harvard University Press.

10. Baake E, Wagner H. 2001 Mutation–selection models solved exactly with methods of statistical mechanics. *Genet. Res.* **78**. (doi:10.1017/S0016672301005110)

11. Bacevic K. 2016 *Cdk2 as a model for studying evolutionary selection and therapeutic responses in proliferating cancer cells*. PhD Thesis. University of Montpelier.

12. Bacevic K *et al.* 2017 Spatial competition constrains resistance to targeted cancer therapy. *Nat. Commun.* **8**. (doi:10.1038/s41467-017-01516-1)

13. Konieczkowski DJ *et al.* 2014 A Melanoma Cell State Distinction Influences Sensitivity to MAPK Pathway Inhibitors. *Cancer Discov.* **4**, 816–827. (doi:10.1158/2159-8290.CD-13-0424)

14. Miyamoto T, Furusawa C, Kaneko K. 2015 Pluripotency, Differentiation, and Reprogramming: A Gene Expression Dynamics Model with Epigenetic Feedback Regulation. *PLOS Comput. Biol.* **11**, e1004476. (doi:10.1371/journal.pcbi.1004476)

15. Melikyan A. 1998 *Generalized Characteristics of First Order PDEs*. Boston, MA: Birkhäuser Boston. (doi:10.1007/978-1-4612-1758-9)

16. Melikyan AA, Ovseevich AI. 1984 Hamiltonian systems with a specified invariant manifold and some of their applications. *J. Appl. Math. Mech.* **48**, 140–145. (doi:10.1016/0021-8928(84)90079-0)

17. Melikyan AA, Ovseevich AI. 2011 Universal surfaces and smooth solutions of Bellman's equations. *Russ. J. Math. Phys.* **18**, 176–182. (doi:10.1134/S1061920811020063)

18. Evans LC. 2014 Envelopes and nonconvex Hamilton–Jacobi equations. *Calc. Var. Partial Differ. Equ.* **50**, 257–282. (doi:10.1007/s00526-013-0635-3)

19. Crandall MG, Ishii H, Lions P-L. 1992 User's guide to viscosity solutions of second order partial differential equations. *Bull. Am. Math. Soc.* **27**, 1–68. (doi:10.1090/S0273-0979-1992-00266-5)
